# PAS domain of flagellar histidine kinase FlrB exhibits novel architecture, and binds Heme as sensory signal in unconventional fashion

**DOI:** 10.1101/2023.06.29.547052

**Authors:** Peeali Mukherjee, Shubhangi Agarwal, Sritapa Basu Mallick, Jhimli Dasgupta

## Abstract

Phosphorylation of the σ^54^-dependent transcription activator FlrC by the sensor histidine kinase FlrB is essential in flagellar synthesis of *Vibrio cholerae*. Despite that, the structure, sensory signal, and mechanistic basis of function of FlrB were elusive. Here we report the crystal structure of the sensory PAS domain of FlrB in functional dimeric state that exhibits a novel architecture. Series of biochemical/biophysical experiments unequivocally established heme as sensory ligand that packs hydrophobically in the ligand binding cleft of FlrB-PAS. Intriguingly, ATP binding to the C-terminal ATP binding (CA) domain assists PAS domain to bind heme, vis-à-vis, heme binding to the PAS facilitates ATP binding to CA; suggesting a synergistic mode of heme and ATP binding to FlrB. We propose that such synergistic binding triggers conformational signaling in FlrB, leading to the downstream flagellar gene transcription. Enhanced swimming motility of *V. cholerae* with increased heme uptake further supports this proposition.

## Introduction

Very often, bacteria respond to the changeable living environments through sensor histidine kinases (HKs), which act together with their cognate response regulators (RRs) to produce necessary downstream responses (Buschiazzo & Trajtenberg, 2019). The two component systems (TCSs) made of HKs and RRs are, therefore, critical for the survival and pathogenicity. Consequently, TCSs are considered as attractive drug targets to combat bacterial infections (Gotoh et al., 2010; Stephenson & Hoch, 2002). The HKs are mostly homodimeric (Buschiazzo & Trajtenberg, 2019) and approximately 83% of HKs contain transmembrane helical regions (Cock & Whitworth, 2007; Ishii & Eguchi, 2021). Those HKs usually possess a variable extracellular sensory domain at the N-terminal of the transmembrane helices, while a cytoplasmic catalytic domain is found at the C-terminus. In that respect, cytosolic HKs are relatively less frequent. HKs contain varieties of sensory domains such as PAS and HAMP which detect broad spectrum chemical and physical stimuli to regulate the activity of the effector domains (Cock & Whitworth, 2007). Usually, ATP binds to the catalytic (CA) domain of HK, autophosphorylates the conserved His residue of the dimerization (DHp) domain. Concomitantly, phosphortransfer occurs to a conserved Asp residue of the cognate RR domain of the partner protein. The phosphorylated RR then triggers downstream responses very often by binding to DNA for transcription regulation of the target gene (Ishii & Eguchi, 2021). Nevertheless, a specific signal must be unequivocally received and interpreted by the sensory domain of the HK, transferred it to the cognate RR for a precise signaling output ensuring high fidelity and no cross-talk.

Gram-negative bacterium *Vibrio cholerae* that causes the diarrheal disease cholera inside the human host is highly motile because of a single, polar sheathed flagellum. *V. cholerae* expresses cholera toxin (Mekalanos, 1983), along with toxin coregulated pilus which are essential for intestinal colonization (R K Taylor, V L Miller, D B Furlong, 1987). Motility, together with colonization at appropriate niche act as the prerequisites for virulence of *V. cholerae*, which in turn are influenced by flagellar synthesis. Similar mechanistic processes for motility, colonization and complete virulence are utilized by other pathogenic bacteria such as *Campylobacter, Helicobacter* species and *Pseudomonas aeruginosa* (Burnham et al., 2020; Eaton et al., 1996; Wassenaar et al., 1993) The transcription of the flagellar genes in *V. cholerae* is organized in a four tired hierarchy (Prouty et al., 2001). The class-I gene product FlrA is responsible to activate σ^54^-dependent transcription of the class-II genes *flrBC* which encode the bacterial enhancer binding protein (bEBP) FlrC and its cognate sensor histidine kinase FlrB (Klose & Mekalanos, 1998). FlrB and FlrC subsequently regulate the expression of class-III genes, encoding basal body hook and the flagellin (Correa et al., 2000; Klose & Mekalanos, 1998).

FlrC is made of N-terminal RR domain followed by the central AAA^+^ ATPase domain and C-terminal DNA binding domain. FlrB phosphorylates a conserved Asp of the RR domain of FlrC (Correa & Klose, 2005b). The structures of the AAA^+^ domain of FlrC (FlrC^C^) (PDB codes: 4QHS, 4QHT), previously determined by us, revealed that FlrC spontaneously forms heptamer without nucleotide dependent subunit remodelling (Dey et al., 2015). Moreover, FlrC atypically binds enhancer elements of *flaA* and *flgK* promoters that are located downstream of σ^54^-binding and transcriptional start sites (Correa & Klose, 2005a). We have recently delineated that high concentration of ubiquitous second messenger c-di-GMP represses ATPase activity of FlrC^C^ by destabilizing its heptameric assembly (Chakraborty et al., 2020). Severe colonization defects of a *V. cholerae* strain were observed upon locking FlrC into either of phosphorylated or unphosphorylated state which implies that subtle regulations of phosphorylation of FlrC by the sensory HK FlrB is required for proper colonization of *V. cholerae* (Correa et al., 2000).

Regardless of the immense importance of the HK FlrB in flagellar gene transcription as well as in motility and colonization of *V. cholerae*, its structure, sensory signal and molecular basis of functions were elusive yet. FlrB of *V. cholerae* was predicted before to be cytosolic with a N-terminal PAS l i ke domain, central histidine kinase phosphortransfer/dimerization domain (DHp) and C-terminal catalytic (CA) domain (Mistry et al., 2021). Autophosphorylation of cytosolic FlrB on a conserved His residue in DHp, followed by phosphotransfer to the RR domain of FlrC was demonstrated previously (Correa et al., 2000).

PAS domains are well known for sequence/functional diversities, and for binding remarkably diverse array of cofactors or ligands. Despite that, PAS domains largely exhibit a conserved core made of five β-strands surrounded by flanking α-helices. Here we report yet unknown crystal structure (at 2.27 Å) of the N-terminal cytosolic PAS domain of *V. cholerae* FlrB (residues 7-123) in its functional dimeric state. Structural comparisons with the reported PAS or PAS like domains depicted a novel architecture of FlrB-PAS with a distinguishingly small globular region. Through systematic docking, and various biochemical and biophysical studies on different FlrB constructs and mutants we have identified heme as the hitherto unknown sensory ligand of FlrB that binds at the globular region of the PAS domain in an unorthodox fashion. Molecular mechanisms underlying the protagonist role of heme in flagellar synthesis are addressed here.

## Results

### Preparation of FlrB constructs

Domain organizations of *V. cholerae* FlrB (Accession code: A0A0H3AMQ9; 1-351 amino acids (aa)), as predicted by SMART, were as follows: PAS domain (aa 21-84), followed by kinase and dimerization (DHp) domain (aa 125-191) and C-terminal ATPase or catalytic (CA) domain (aa 234-340). To start with, we cloned and purified full-length FlrB (FlrB-Fl, aa 1-351) that had severe solubility problem, and it was not suitable for structural and functional studies. However, truncation of a few floppy residues at both the termini yielded a construct FlrB-Fl^7-340^ (7-340 aa), that contains all the domains. FlrB-Fl^7-340^ is soluble, functional, and amenable for structural and functional studies. Existing literature suggests that PAS domains are usually made of ∼100 amino acids (Henry & Crosson, 2011). Since PAS domains share very little primary sequence identity (∼20%), prediction of FlrB-PAS domain based upon sequence resemblance/Pfam data would be erroneous. We, therefore, critically analysed the available PAS domain structures, and prepared two constructs namely FlrB-PAS^7-108^ (aa 7-108) and FlrB-PAS^7-123^ (aa 7-123). All the constructs were overexpressed well in BL21 (DE3) and purified by Ni-NTA affinity chromatography as 6×His-tagged proteins (Fig. S1).

### Structure determination of FlrB-PAS^7-123^

Single crystals of FlrB-PAS^7-108^ and FlrB-PAS^7-123^ were grown by hanging drop vapor diffusion method. FlrB-PAS^7-108^ crystals diffracted poorly up to 4.2 Å. However, FlrB-PAS^7-123^ crystals diffracted well, and a dataset was collected up to 2.27 Å (Table 1). The crystals of FlrB-PAS^7-123^ belonged to the space group C2 with unit-cell parameters a=212.64 Å, b=49.56 Å, c=146.83 Å, β=92.53° (Table 1). Calculation of Matthews coefficient indicated the most probable presence of twelve molecules (considering the molecular weight of 14.8 kDa) in the asymmetric unit, with a V_M_ (Matthews, 1968) of 2.18 Å Da^-1^ and the solvent content of 43%.

**Table 1:** Data-collection and refinement parameters (PDB ID 7YRT)

|  |  |
| --- | --- |
| Protein | FlrB-PAS<sup>7-123</sup> |
| Diffraction source | ESRF BEAMLINE ID23-1 |
| Wavelength (Å) | 0.97242 |
| Temperature (K) | 100 |
| Detector | DECTRIS EIGER2 X 16M |
| Crystal-to-detector distance (mm) | 321.70 |
| Rotation range per image (°) | 1 |
| Total rotation range (°) | 900 |
| Exposure time per image (s) | 0.01 |
| Space group | C2 |
| Unit-cell parameters | a=212.644, b=49.556, c=146.830,<br>β=92.53 |
| Mosaicity (°) | 0.14 |
| Resolution range (Å) | 48.26-2.27 |
| Total No. of reflection | 178295 (17953) |
| No. of unique reflection | 69328 (6819) |
| Completeness (%) | 97.7 (98.6) |
| Multiplicity | 2.6 |
| CC<sub>1/2</sub> | 99.7 (52.6) |
| ⟨I/σ(I)⟩ | 9.1 (1.2) |
| R<sub>merge</sub> (%) | 4.6 (79.7) |
| R<sub>pim</sub> (%) | 4.6 (79.2) |
| Overall B factor from Wilson plot (Å<sup>2</sup>) | 61.8 |
| **Refinement** |  |
| Resolution range (Å) | 48.26-2.27 |
| No. of reflections used in refinement | 69160 |
| R<sub>work</sub>/R<sub>free</sub> | 18.49/25.07 |
| No. of atoms |  |
| Protein | 10957 |
| Solvent | 94 |
| Ligand | 0 |
| RMS bond lengths (Å) | 1.062 |
| RMS bond angles (°) | 0.008 |
| Ramachandran plot |  |
| Favored (%) | 96.39 |
| Allowed (%) | 3.61 |

Since all the PAS domains, irrespective of their low sequence identities, form conserved globular core region with five β stands of varying length, we tried Molecular replacement (MR) calculations using PHENIX (Adams et al., 2010) with different PAS domain structures (of PDB codes 1DRM, 1Y28, 3VOL, 6CEQ, 1V9Z, 2R78, 3LYX, 4GCZ, 1LL8 etc). However, MR calculations did not produce any satisfactory solution indicating that an alternative method is to be availed to solve the structure. Finally, the monomeric model of FlrB-PAS^7-123^, predicted by AlphaFold2 (Mirdita et al., 2022), produced clear-cut MR solution with R_work_ of 37.4% and R_free_ of 41.7% for twelve monomers arranged in the form of six dimers in the asymmetric unit. The structure of FlrB-PAS^7-123^ were further refined to R_work_ of 19.3% and R_free_ of 25.0%. As correctly pointed out by McCoy et al (2022) (McCoy et al., 2022), the structures that were the most challenging to solve with the AlphaFold2 models contained extended helices, especially with kinks. Although the crystal structure of FlrB-PAS^7-123^ and the model predicted by AlphaFold2 are similar with respect to the globular region, the N-terminal helix of the crystal structure is significantly different from the predicted model (Fig. S2).

### Structure of FlrB-PAS^7-123^, and its interactions at the dimeric interface

In general, the globular regions of the PAS domains are found to be of ∼100 amino acid residues. Our structural study delineated that the globular region of FlrB-PAS^7-123^ is small and is made of only 78 amino acids (aa 24-102). The β-sheet made of five β-strands, Aβ, Bβ, Eβ, Fβ, and Gβ constitutes the core part of the globular region (Fig. 1A, B). Helices Cα, Dα, along with the extended loops Cα-Dα and Dα-Eβ flank the β-sheet (Fig. 1B, C). The N-terminal and C-terminal helices are nomenclatured as Aα′ and Hα respectively (Fig. 1B, C).

**Figure 1:**
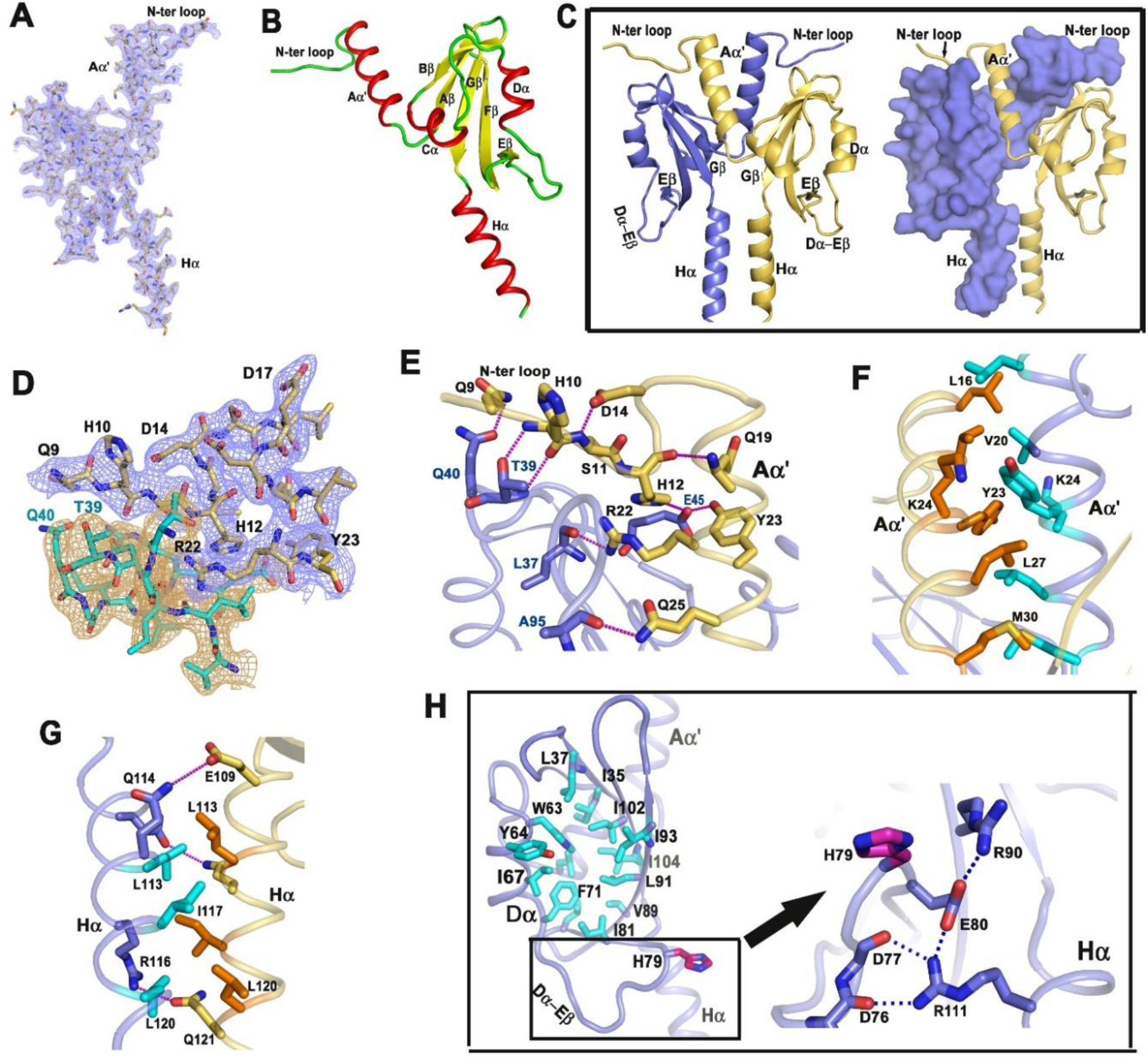
Crystal structure and inter-subunit interactions of FlrB-PAS^7-123^. (A) 2F_O_-F_C_ electron density (contoured at 1σ) overlaid on the refined model of FlrB-PAS^7-123^. (B) Cartoon diagram of FlrB-PAS^7-123^. (C) Dimer of FlrB-PAS^7-123^. Interactions at the dimeric interface are shown in cartoon and surface model. (D) 2F_O_-F_C_ electron density (contoured at 1σ) around the N-terminal loop that interacts with the globular region of the *trans*-subunit. (E) Detailed interactions between N-terminal loop of one monomer and the globular region of the *trans*-subunit. (F) Hydrophobic packing between Aα′ helices of the two monomers. (G) Hydrophobic and polar interactions between the two C-terminal Hα helices. (H) The left panel shows hydrophobic pocket in the globular region of FlrB-PAS^7-123^. Right panel is the zoomed view of Dα-Eβ loop showing polar/salt bridge interactions between the loop with neighbouring β strand and Hα.

Dimerization of FlrB-PAS^7-123^ occurs primarily due to the inter-subunit interactions between the N-terminal Aα′ helices, between the C-terminal Hα helices, and between Aα′ of one subunit with the globular region of the other (Fig. 1C). Contact surface areas between the subunits, as calculated by PDBePISA, is 1995 Å^2^ where free energy of stabilization upon dimerization is ∼ ̶ 30 kcal/mole. Polar interactions of the extended N-terminal loop with the globular domain of the other subunit evidently maximize inter-subunit interactions (Fig. 1C-E). N-terminal Aα′ helices of the two subunits packed together through hydrophobic interactions (Fig. 1F), while both polar and hydrophobic interactions were observed between C-terminal Hα helices (Fig. 1G). One side of Aα′ was involved in inter-helix interactions and the other side of it was found to interact with the globular region of the *trans*-subunit. The core of the globular region, which is presumed to be the ligand binding pocket of FlrB-PAS, is made of hydrophobic residues namely Leu37, Trp63, Phe 71, Leu91, Ile93, Ile104 etc. (Fig. 1H). Salt bridge interactions are observed between Asp76 and Glu80 of the Dα-Eβ loop and Arg111 of the C-terminal helix Hα (Fig. 1H).

### PAS domain of FlrB possesses novel architecture

Structural comparison performed in the DALI server (http://ekhidna2.biocenter.helsinki.fi/dali/), using the refined structure of FlrB-PAS^7-123^ as search model, identified only few neighbours but with high Z-scores (and high RMSD values) (Table S1). Interestingly, some of these identified neighbours were already tested as MR search models and failed (Table S1). Aforesaid facts indicated that N-terminal PAS domain of FlrB possesses a novel architecture. Structures of PAS domains involved in binding p-coumaric acid, 3,5-dimethyl-pyrazine-2-ol (DPO), FAD or heme revealed that they all have a ligand binding cleft made of the core antiparallel β-sheet and several flanking α-helices (Möglich et al., 2009) (Fig. 2A-E). A nomenclature of the alpha helices and beta sheets that constitute the globular PAS core has been defined by Stuffle et al (2021) (Stuffle et al., 2021). The antiparallel β-sheet, made of the strands Aβ, Bβ, Gβ, Hβ, and Iβ is considered to be the most conserved region. But the orientation, length, and number of flanking α-helices, designated as Cα, Dα, Eα, and Fα, and their connecting loops differ substantially (Fig. 2A-D). The adjoining N-terminal helix associated with the PAS core is named as (Aα′) while the C-terminal helices that pack against the β-sheets are named as Jα. Structures of the PAS domains involved in FAD and heme binding (Fig. 2B, D) indicate that five stranded β-sheet forms the base of the ligand binding cleft where the flanking Dα, Eα, and Fα act as a cover. The structure of FlrB-PAS^7-123^ is non-superimposable with any of these PAS structures. An apparent resemblance in the β-sheet region has been observed between the PAS domain of FlrB and the transcription factor TyrR although the RMSD value of superposition (4.0 Å for 74 atoms) was exorbitantly high (Fig. 2E).

**Figure 2:**
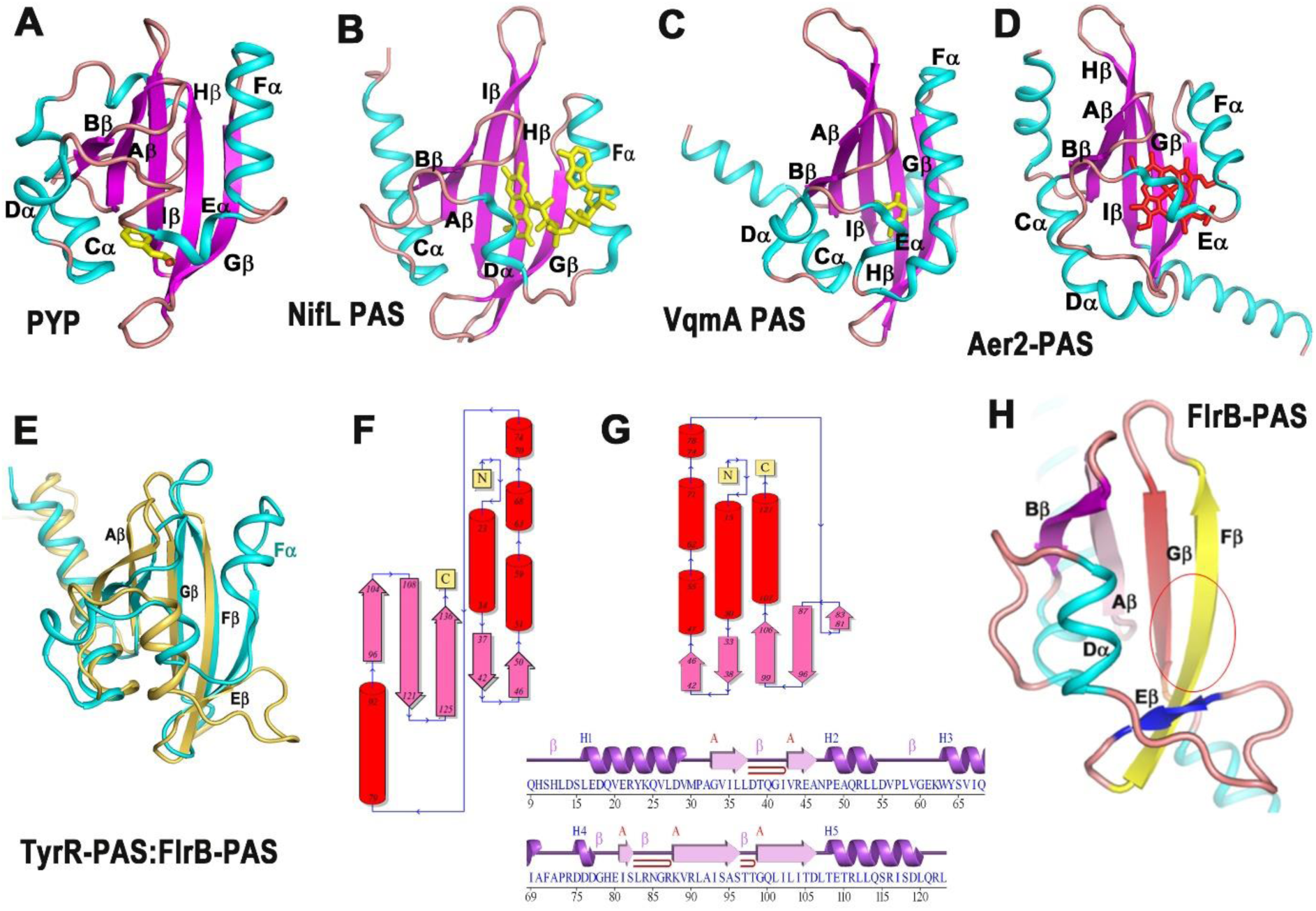
Novel fold of the globular domain of FlrB-PAS^7-123^. (A) Crystal structures of photoactive yellow protein (PYP) from *Halorhodospira halophila* that resembles PAS with its cofactor p-coumaric acid (PDB code: 1NWZ), (B) PAS domain of NifL that binds FAD (PDB code: 2GJ3), (C) PAS domain of *V. cholerae* VqmA bound to the ligand 3,5-dimethyl-pyrazine-2-ol (DPO) (PDB: 6UGL). (D) PAS domain of Aer2 from *P. aeruginosa* that binds heme (PDB code: 3V0L). (E) Superposition of the PAS domain of gene regulatory protein TyrR of *E. coli* (cyan) on FlrB-PAS^7-123^ (gold) shows drastic difference in architecture of two PAS domains. (F, G) Topologies of the common PAS domain (F), and of FlrB-PAS (G), generated by PDBsum further establish novel architecture of FlrB-PAS. (H) Probable ligand binding pocket of FlrB-PAS^7-123^ marked by red circle.

The structural uniqueness of FlrB-PAS^7-123^ resides in its topology. A comparison of the topology between common PAS fold and that of FlrB-PAS (Fig. 2F,G) showed that the globular part of FlrB-PAS^7-123^ structure is substantially smaller compared to the common PAS fold (Fig. 2F,G). Importantly, Eβ of FlrB-PAS^7-123^, that corresponds to the Gβ of the other PAS domains, is significantly smaller in size (Fig. 2E-H). Additionally, FlrB-PAS^7-123^ is devoid of two flanking helices (Fig. 2G, H). Generally, ligand packs between the β-sheet and Fα (Fig. 2A-D), and that pivotal helix Fα does not exist in FlrB-PAS^7-123^ (Fig. 2G, H). Instead, FlrB-PAS^7-123^ contains an extended loop between Dα and Eβ (Fig. 2H). Altogether, the globular region of FlrB-PAS^7-123^ exhibits a novel architecture with a relatively open ligand binding pocket (Fig. 2).

### Identification of the sensory ligand of FlrB-PAS

Deciphering yet unknown sensory role of FlrB-PAS was the most challenging. Being a cytosolic HK, it is logical to speculate that the PAS domain of FlrB acts as an intracellular sensor, and binds the signalling molecule that enters in to the cytosol. Sensory domains of the histidine kinases such as FixL of *Bradyrhizobium japonicum,* Aer2 from *Pseudomonas aeruginosa* or HrrSA and ChrSA of Gram-positive soil bacterium *Corynebacterium glutamicum* were seen to bind heme in a hydrophobic pocket (Garcia et al., 2017; Keppel et al., 2018). Histidine, tyrosine or His/Tyr are well known to be as the iron-coordinating residues (Garcia et al., 2017; Keppel et al., 2018). However, several studies recently depicted binding of heme solely through the hydrophobic packing without contributions from His or Tyr (Gao et al., 2018; Kühl et al., 2011; Rao et al., 2011; Tan et al., 2013). Structure of FlrB-PAS^7-123^ revealed that the globular region possesses Tyr64 and His79 along with various hydrophobic residues, distributed quite proportionally all over the ligand binding pocket, raising possibility of heme binding (Fig. 1H). Nevertheless, we initiated our ligand search with p-coumaric acid, 3,5-dimethyl-pyrazine-2-ol (DPO), FAD, and heme.

#### Autodock and ProBis indicated heme binding in FlrB-PAS^7-123^

Docking experiments were performed with Autodock Vina using FlrB-PAS^7-123^ coordinates. The parameters were kept as center_x = 6.101, center_y = -1.926, center_z = 61.53. Size of blind box was adjusted to size_x = 52, size_y = 72, size_z = 52. Results of Autodock suggested that FlrB-PAS^7-123^ has highest binding affinity of -7.2 kcal/mol towards heme with rmsd l.b as zero, followed by FAD, and negligible binding probability with p-coumaric acid and DPO.

Recently, ProBis has emerged as an efficient server for ligand detection (Konc & Janežič, 2014). Interestingly, an unbiased ligand search with the coordinates of FlrB-PAS^7-123^ (PDB code: 7YRT) undisputedly identified protoporphyrin IX as its specific ligand (Fig. S3).

#### ITC experiment supported heme binding by FlrB-PAS^7-123^

Although, binding of p-coumaric acid, 3,5-dimethyl-pyrazine-2-ol (DPO), FAD with FlrB-PAS^7-123^ were less probable compared to heme, we have tested all four ligands for FlrB-PAS^7-123^ binding through ITC experiments. Binding parameters of Heme b or hemin (PubChem SID: 329815257) to FlrB-PAS^7-123^ were characterized by using reverse titration technique (Vu et al., 2013). Hemin was freshly prepared in DMSO and diluted with buffer prior to the experiment. FlrB-PAS^7-123^ was also equilibrated with equal percentage of DMSO. Heat of dilution was subtracted from corresponding FlrB-PAS^7-123^ hemin binding before curve fitting. Titration between FlrB-PAS^7-123^ and hemin was exothermic and early saturation was achieved (Fig. 3A). Binding isotherm were fitted using the one site binding model using Origin8.5 software. Titration confirmed binding between hemin and FlrB-PAS^7-123^ with dissociation constant K_d_ of 7.72 ± 0.16 µM (Fig. 3A). However, no considerable binding was observed for p-coumaric acid and DPO, while an endothermic graph was observed with FAD with no saturation. Nonetheless, none of these plots produced meaningful K_d_ value after buffer subtraction (Fig. S4).

**Figure 3:**
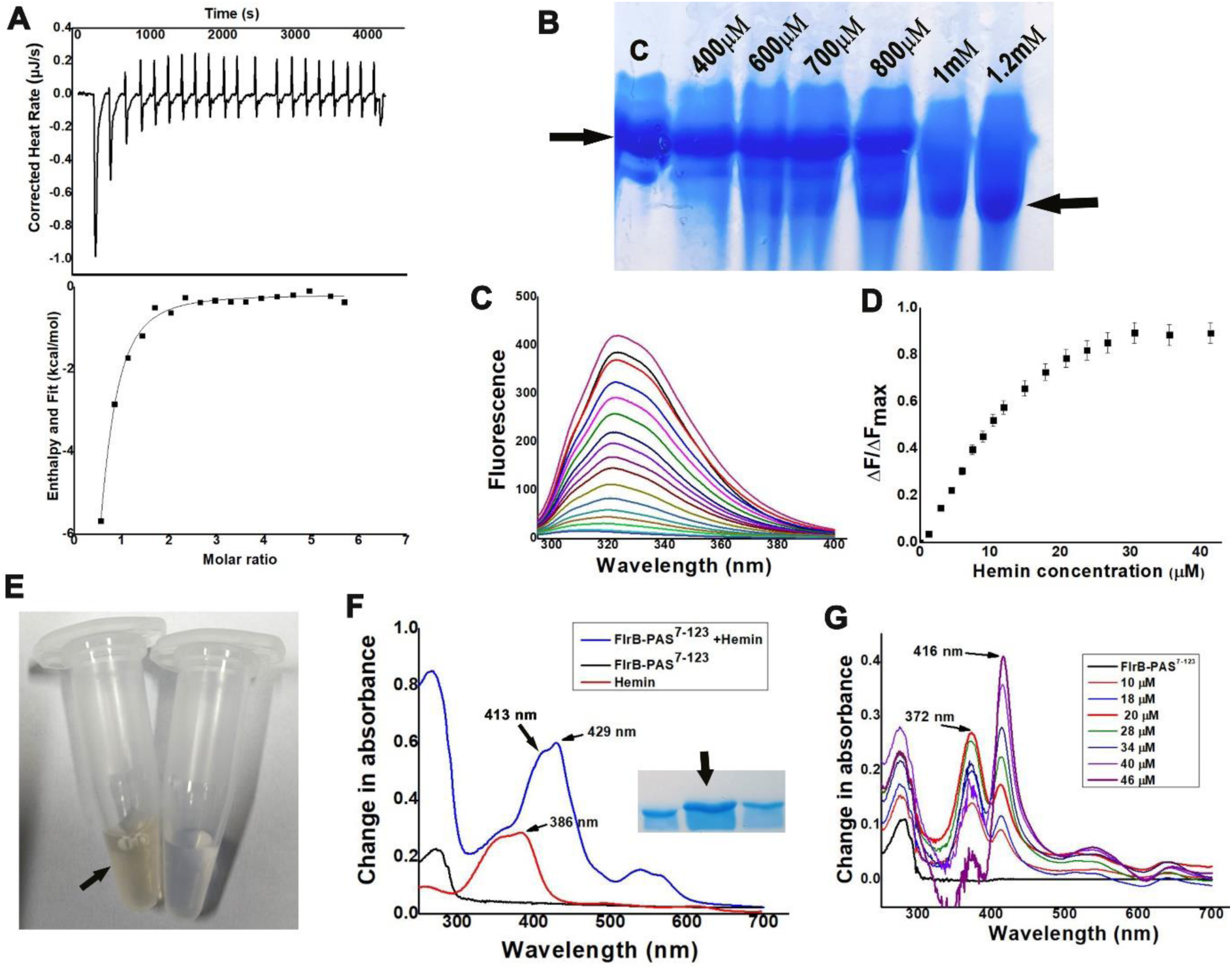
Hemin binding to FlrB-PAS^7-123^. (A) Hemin binding was observed through Isothermal Titration Calorimetry with K_d_ of 7.72 ± 0.16 µM. The thermogram shows heat change along with lower panel of integrated isotherms. (B) 12% native PAGE of FlrB-PAS^7-123^ incubated with increasing concentration of hemin (mentioned over the respective lanes). Lane ‘C’ stands for the protein FlrB-PAS^7-123^ of 200 μM without hemin. (C, D) Changes in fluorescence quenching upon hemin binding with FlrB-PAS^7-123^ where λ_exc_=280 nm and λ_em_=295−400 nm. Graph of ΔF/ΔF_max_ vs hemin concentration produced K_d_ of 8.5±0.02 μM. (E) FlrB-PAS^7-123^ purified after addition of hemin in cell lysate is yellowish in colour (left side), while the right-side aliquot (transparent) is FlrB-PAS^7-123^ purified without hemin. Hemin was added to the lysate after sonication and purified as per protocol of FlrB-PAS^7-123^ without hemin. (F) Eluted fraction of FlrB-PAS^7-123^ with hemin was subjected to UV-vis spectral scan. Soret band observed at 413 nm and 429 nm confirmed hemin binding. Spectral lines of protein alone is in black, hemin alone is in red and of hemin treated protein is in blue. Fraction used for this experiment is shown as inset. (G) FlrB-PAS^7-123^ was titrated with increasing concentration of hemin (no protein in reference cell) and scanned after incremental addition of hemin. Change in absorbance versus wavelength was plotted after subtracting blank (hemin titration in the absence of protein). Experiments were performed in triplicates and error bar is given after calculating ± (SD). Hemin of 3 mM was used for fluorescence quenching and UV-vis spectroscopic experiments.

#### Migration pattern in native PAGE demonstrates hemin binding with FlrB-PAS^7-123^

We performed native PAGE to qualitatively inspect hemin binding with FlrB-PAS^7-123^. Experiments were carried out at pH 8.0, where 200 μM of FlrB-PAS^7-123^ was incubated with gradually increasing concentrations (0.4 mM to 1.2 mM) of hemin. Inspection of Coomassie blue-stained gel clearly indicate two different migration pattern one for free FlrB-PAS^7-123^ and the other for hemin bound FlrB-PAS^7-123^, where the latter migrate faster (Fig. 3B). Distinct band for hemin bound FlrB-PAS^7-123^ started to appear with 600 μM of hemin and maximized at 1.2 mM of hemin (Fig. 3B). Faster migration of the hemin bound form compared to the apo form could be accounted for the propionate groups of the hemin molecules. Notably, faster migrations upon hemin binding were demonstrated before for HutB of *V. cholerae* (Agarwal et al., 2017), or HbpA of Haemophilus influenza (Vergauwen et al., 2010).

#### Fluorescence quenching experiments confirmed hemin binding with FlrB-PAS^7-123^

To monitor probable interactions between hemin and FlrB-PAS^7-123^, we have conducted fluorescence quenching studies. FlrB-PAS^7-123^ showed substantial quenching upon addition of hemin, with a K_d_ value of 8.5 ± 0.02 μM which is indicative of efficient hemin binding (Fig. 3C, D). Interestingly, FlrB-PAS^7-123^ hosts a solitaire Tryptophan (Trp63) and Tyr64 in its globular region. Therefore, our quenching studies provide a solid clue that the hemin binding site resides at the globular region and Trp63/Tyr64 are within the Forster distance of the potential ligand binding site. The change in the intrinsic fluorescence of FlrB-PAS^7-123^ upon hemin binding were monitored (in triplicate) at excitation wavelength (λ_exc_) of 280 nm and emission wavelength (λ_em_) of 295−400 nm.

As a positive control, we have tested binding of the hemin with BSA through fluorescence quenching (using same parameters stated above), as BSA is well known to bind heme (Flint & Stintzi, 2015). Significant fluorescence quenching of BSA produced K_d_ value of 1.78 ± 0.02 μM (Fig. S5).

#### Shift of Soret band of hemin was observed upon binding with FlrB-PAS^7-123^

The affinity of heme for FlrB-PAS^7-123^ was further checked by adding hemin to the cell lysate before purification of FlrB-PAS^7-123^. Yellowish colour of FlrB-PAS^7-123^ after purification through Ni-NTA affinity chromatography indicated hemin binding with FlrB-PAS^7-123^ (Fig. 3E). Hemin binding with FlrB-PAS^7-123^ was further confirmed by UV-Vis spectroscopic experiments. The absorption spectrum of hemin shows a peak in the Soret band region around 386 nm (Fig. 3F). Bathochromic shift of the Soret band implies stable protein-heme interaction (Karnaukhova et al., 2014). Apo FlrB-PAS^7-123^ does not show any absorption in the Soret region (Fig. 3F). But FlrB-PAS^7-123^ purified in the presence of hemin has shown a clear-cut bathochromic shift of the Soret band with λ_max_ at 413 nm followed by 429 nm when the spectral range was measured between 250 – 700 nm (Fig. 3F). Hemin was incrementally titrated to FlrB-PAS^7-123^ to see the effect of heme:protein ratio on Soret shift. Upon titration, a blue shift of hemin Soret band was observed with λ_max_ at 372 nm followed by a red shift with λ_max_ at 416 nm with respect to hemin peak alone at 385 nm (Fig. 3F, G) with gradual increase in hemin concentration. Such split Soret bands are indicative of hexa- and penta coordination of heme with the protein (Mattle et al., 2010; Wißbrock et al., 2019). The exact location of the shifted Soret bands, however, depends on the mode of binding and immediate surroundings of the hemin molecule (Luthra et al., 2011). Saturation of the blue shifted peak was obtained with hemin concentration of 20 µM (where FlrB-PAS^7-123^ concentration was 10 µM) (Fig. 3G). After that, an increment in intensity of the red shifted peak was observed with gradual decrease in intensity of the blue shifted peak upon further addition of hemin (Fig. 3G). The isosbestic point at 404 nm implied a two-state conversion. While blue shifted Soret peak was indicative of the initial high energy interaction between FlrB-PAS^7-123^ and hemin, red shift to 416 nm underscored more stable protein-hemin complex.

### FlrB-PAS exists as dimer in solution in free and heme bound states

In view of structural and functional observations, we have tested the presence of the dimeric assembly of FlrB-PAS^7-123^ in solution by size exclusion chromatography (SEC) (Fig. 4A, S6). Similar experiments were carried out with FlrB-PAS^7-108^ since this construct is devoid of the C-terminal helix, Hα (Fig. 4B, S6). Both the constructs were observed in the dimeric states (Fig. 4A, B). Higher hydrodynamic radius is often observed because of elongated molecular architecture and/or oligomeric sates. Relatively higher hydrodynamic radius of FlrB-PAS^7-123^, calculated from the elution volume, might be attributed to elongated C-terminal helical region (Fig. S6). Qualitative SEC experiments, for both the constructs, in three different protein concentrations (500 μM, 250 μM and 100 μM) exhibited negligible shifts in peak positions demonstrating consistent presence of dimeric states irrespective of dilutions (Fig. 4A, B). Chemical crosslinking experiments of FlrB-PAS^7-123^ and FlrB-PAS^7-108^ using Ethylene glycol bis-succinimidyl succinate (EGS) as crosslinker is done with successively increasing concentrations (Fig. 4C, D). Presence of dimeric (∼30 kDa) form of FlrB-PAS^7-123^ with increasing EGS concentration is evident from the migration pattern of the crosslinked product run in 12% SDS-PAGE gel (Fig. 4C). Sustainable dimer of FlrB-PAS^7-108^ were also observed both in SEC and crosslinking experiments (Fig. 4B-D). Retention of dimeric state of FlrB-PAS^7-108^, that lacks C-terminal helix Hα, highlighted the importance of inter-subunit interactions between the N-terminal loop and globular region in the stability of dimeric structure (Fig. 1C, 4B, D).

**Figure 4:**
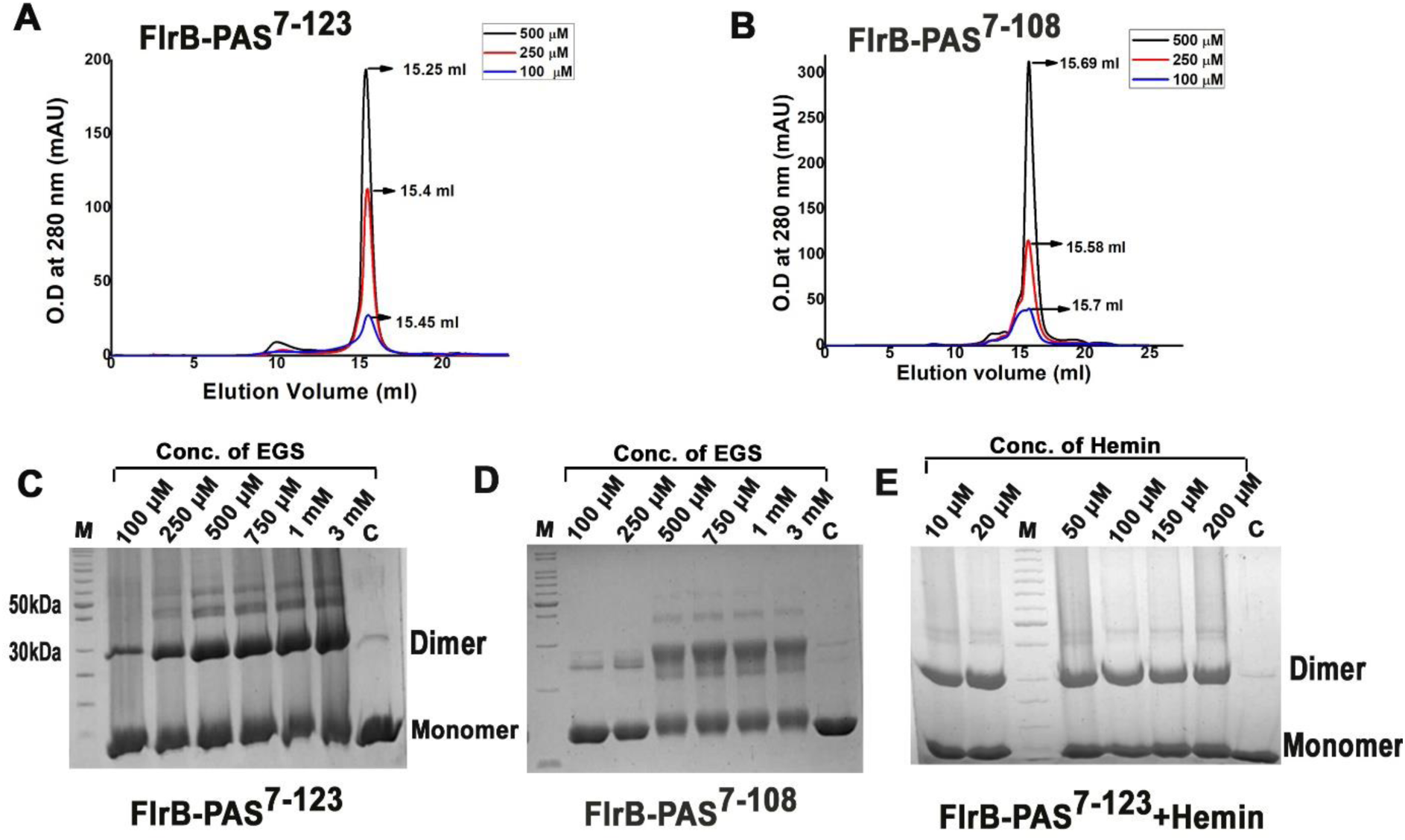
Stable dimerization of the PAS domain of FlrB. (A) Size exclusion chromatography profile of FlrB-PAS^7-123^ showed dimer. The chromatogram profile shows protein of 100 µM, 250 µM and 500 µM concentration where right shift of the peak was visible upon dilution. (B) Size exclusion chromatography of FlrB-PAS^7-108^ also showed dimer. Similar protein concentrations were used here. (C) FlrB-PAS^7-123^ dimerization was further investigated through crosslinking experiment. 2mg/ml of FlrB-PAS^7-123^ was incubated with increasing concentration of crosslinker EGS for 30 mins. Appropriate dimer band is visible near 30 kDa. Intensity of this band is increasing upon increasing concentration of EGS (concentration of EGS is mentioned over the respective lane). (D) PAS^7-108^ construct was crosslinked with increasing concentration of EGS. Intense dimer band is visible from 500 µM to 3 mM of EGS. (E) No effect of hemin was observed on FlrB-PAS^7-123^ dimerization during crosslinking of hemin incubated protein. Hemin concentration is labelled over the respective lanes. All samples were crosslinked with 500 µM EGS for 30 mins. Saturation of dimer is visible from 10 µM of hemin with decreased nonspecific higher molecular weight protein bands. 12% SDS PAGE was used for the experiment. In this Figure, lane ‘C’ stands for FlrB-PAS^7-123^ in (C, E) and FlrB-PAS^7-108^ in (D), in absence of EGS/hemin.

Effect of heme on dimerization was investigated through similar crosslinking experiments where FlrB-PAS^7-123^ was pre-incubated with increasing concentrations of hemin from 10 μM to 200 μM (Fig. 4E). Since hemin was dissolved in DMSO, control cross-linking experiments were carried out after incubation of the protein with equivalent concentration of DMSO without hemin. However, no effect of DMSO was observed there (data not shown). Hemin binding did not hinder dimerization, rather non-specific high molecular weight bands were diminished in the presence of hemin (Fig. 4E). This result indicated that dimerization interface and heme binding regions are distinct.

### Identification of the mode of heme binding through mutational analysis

Soaking of FlrB-PAS^7-123^ crystals with hemin did not produce any hemin bound crystal. In fact, attempt to crystallize hemin incubated FlrB-PAS^7-123^ yielded the crystals of the apo form leaving hemin in the mother liquor (Fig. S7A). The reasons behind such results were hidden in the crystal packing of FlrB-PAS^7-123^ (Fig. S7B, C). Each monomer of a FlrB-PAS^7-123^ dimer packs with the neighbouring dimer through hydrophobic and polar interactions that involve the ligand binding pocket of the globular region (Fig. S7B, C). According to PDBePISA, the stability gained due to such crystal packing was significant with Δ*G* of ∼ ̶ 10 kCal for each dimer and of ∼ ̶ 60 kCal for the whole asymmetric unit. Understandably, such packing of FlrB-PAS^7-123^ molecules in the present crystal form is reasonably stable to outcompete the binding of hemin. Although we obtained hemin bound crystals of FlrB-PAS^7-108^ (Fig. S7D), possibly packed in a different space group, low resolutions of the data restricted the structure determination.

We, therefore, decided to determine the mode of hemin binding through mutational analysis. We have observed that heme binding does not interfere with dimerization. Thus, we have investigated the heme binding properties of FlrB-PAS^7-108^, the construct devoid of Hα (Fig. 5A). Fluorescence quenching experiments of FlrB-PAS^7-108^ (λ_exc_=280 nm, λ_em_=295−400 nm) with hemin indicated binding with a K_d_ value of 11 ± 0.1 μM, which is comparable to that of FlrB-PAS^7-123^ (Fig. S8A, 5B, 5D). UV-vis spectroscopy depicted predominant blue shifted Soret peak with λ_max_ at 372 nm, and a negligible red shifted Soret peak at λ_max_ of 413 nm upon titration with FlrB-PAS^7-108^ (Fig. 5C). However, the red shifted Soret band, which was at 416 nm in case of FlrB-PAS^7-123^ was practically absent here (Fig. 5C). Nonetheless, overall results were indicative of hemin binding to the globular region of FlrB-PAS^7-108^.

**Figure 5:**
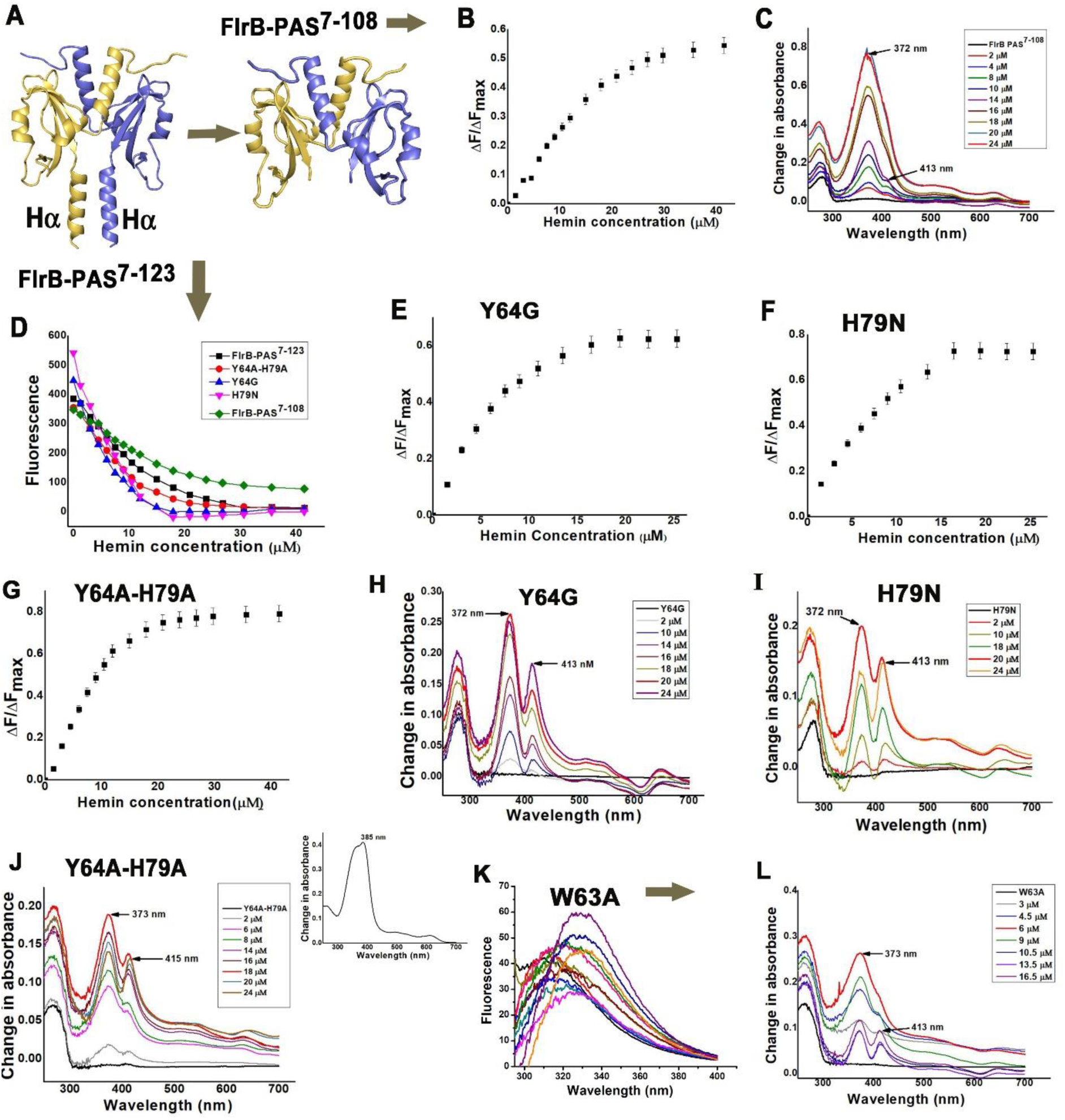
Hemin binding with FlrB-PAS^7-108^. (A) FlrB-PAS^7-108^ model prepared from the coordinates of the crystal structure of FlrB-PAS^7-123^. (B) FlrB-PAS^7-108^ was titrated with increasing concentration of hemin and quenching in fluorescence intensities were measured at λ_exc_=280 nm and emission at λ_em_=295−400 nm. Graph of ΔF/ΔF_max_ vs hemin concentration produced K_d_ of 11± 0.1 μM. (C) UV-vis spectroscopy analysis of heme titration to FlrB-PAS^7-108^ showed predominantly blue shift of Soret band. (D) Decrease in fluorescence for FlrB-PAS^7-108^, FlrB-PAS^7-123^ and its mutants Y64G, H79N and Y64A-H79A are plotted against hemin concentration to show saturations. (E, F, G) Plot of ΔF/ΔF_max_ versus hemin concentrations for Y64G, H79N and the double mutant Y64A-H79A produced K_d_ values of 4.28 ± 0.01 µM, 4.9 ± 0.01 µM, and 6.93 ± 0.02 µM respectively. (H, I) Hemin titration showed blue and red shifts of Soret bands to 372 nm and 413 nm for Y64G and H79N respectively. (J) Similar blue and red shifts for the double mutant Y64A-H79A at 373 nm and 415 nm indicated binding with hemin. Soret band of hemin only (inset) showed a peak at 385nm for all experiments. (K) W63A mutant did not show proper initial fluorescence intensity. Quenching was noisy and was not suitable to calculate K_d_ value. (L) UV-vis spectroscopic analysis showed blue shift of Soret band to 373 nm with negligible red shift. 3 mM hemin stock was used for all fluorescence quenching and UV-vis spectral wavelength scan experiments.

We prepared three point-mutants W63A, Y64G, H79N, and a double mutant Y64A-H79A on FlrB-PAS^7-123^ for hemin binding studies. The mutants were overexpressed and purified (Fig. S1B) as per protocol mentioned for FlrB-PAS^7-123^. Fluorescence quenching and UV-vis spectroscopic studies were carried out with the mutants using the protocols mentioned above. Similar to FlrB-PAS^7-123^, significant decrease in fluorescence upon titration with hemin were observed for Y64G, H79N and Y64A-H79A (Fig. S8B-D, 5D). Corresponding K_d_ values for Y64G, H79N and Y64A-H79A were 4.28 ± 0.01 µM, 4.9 ± 0.01 µM, and 6.93 ± 0.02 µM respectively indicating comparable binding affinity of hemin as observed for FlrB-PAS^7-123^ (Fig. 5E-G).

Beside this, hemin was incrementally titrated to 10 μM of each mutant at pH 8.0, and the spectral range between 250 –700 nm was monitored (Fig 5H-J). While blue shifted Soret bands were observed at 372 nm for Y64G, H79N and Y64A-H79A, red shifted Soret bands were at 413 nm for Y64G, H79N, and at 415 nm in case of Y64A-H79A (Fig 5H-J).

Drastic reduction in the initial fluorescence of the mutants W63A indicated that the overall fluorescence of FlrB is primarily contributed by Trp63 (Fig. 5K). Although quenching was observed upon addition of hemin to W63A, the noisiness of the data didn’t allow us to calculate the K_d_ value. During spectroscopic analysis of W63A blue shifted Soret band was primarily observed at 373 nm with a weak red shifted band at 413 nm (Fig. 5L).

### Synergistic mode of heme and ATP binding to FlrB-Fl^7-340^

We investigated hemin binding to FlrB-Fl^7-340^ as well. Similar to FlrB-PAS^7-123^, addition of hemin in the cell lysate during purification of FlrB-Fl^7-340^ produced coloured protein, and bathochromic shift of the Soret band to 414 nm confirmed hemin binding with FlrB-Fl^7-340^ (Fig. 6A,B). Interestingly, titration of FlrB-Fl^7-340^ (concentration 10 μM) with gradual increment of hemin exhibited only a red shifted Soret band with λ_max_ at 415 nm that points towards stable protein-hemin interactions (Fig. 6C). Intensity of the peak was increasing with increase in hemin:protein ratio, although saturation was not achieved even with 34 μM of hemin. Since FlrB-Fl^7-340^ contains ATP binding domain too, we checked probable influence of AMP.PNP (non-hydrolysable ATP analog) binding in the CA domain on hemin binding to the PAS domain. FlrB-Fl^7-340^, pre-incubated with AMP.PNP, showed a red shift peaking at 416 nm. However, in this case, a trough at 360-390 nm accompanied the peak at 416 nm, and saturation was obtained very quickly only with 4 μM of hemin (Fig. 6D). Further addition of hemin (to 34 μM) caused gradual red shift with a peak at 420 nm (Fig. 6D). This result indicated that binding of AMP.PNP in the CA domain of FlrB-Fl^7-340^ plausibly assists hemin binding in the PAS domain, and an equilibrium of this dual ligand bound state is obtained easily.

**Figure 6:**
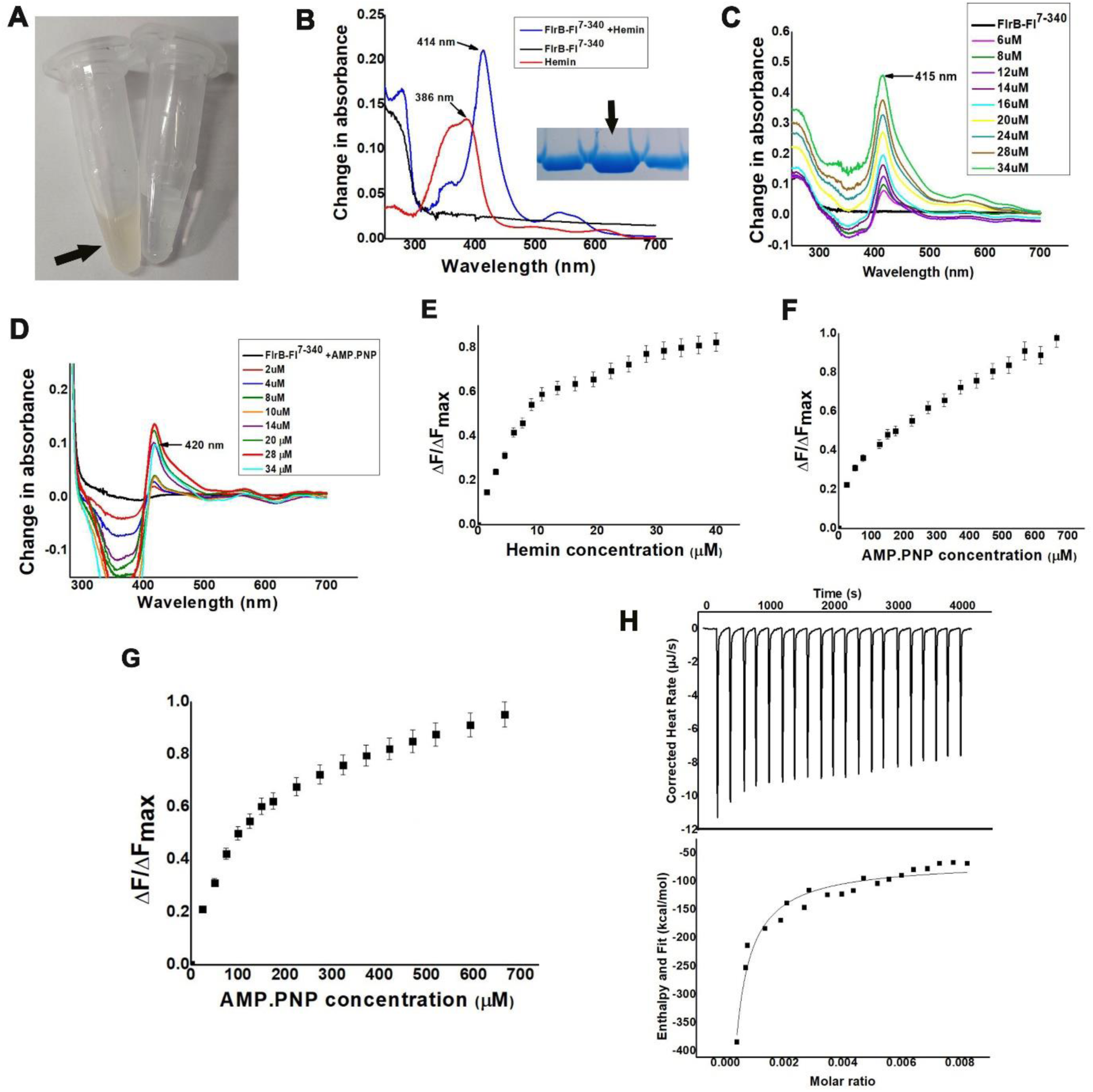
Synergistic mode of hemin and AMP.PNP binding to FlrB-Fl^7-340^. (A) FlrB-Fl^7-340^ purified in presence of hemin in lysate showed eluted protein of yellowish colour. Left aliquot is fraction of FlrB-Fl^7-340^ purified in the presence of hemin and right aliquot is purified without hemin. (B) Red shift of Soret band peaking at 414 nm confirmed hemin binding with FlrB-Fl^7-340^ purified in the presence of hemin. Spectral lines of protein alone is in black, hemin alone is in red and of hemin treated protein is in blue. Fraction used for this experiment is shown as inset. (C) Hemin titration to FlrB-Fl^7-340^ produced red shifted Soret band with λ_max_ at 415 nm. (D) FlrB-Fl^7-340^ preincubated with AMP.PNP showed red shifted Soret band at λ_max_ at 416 nm with only 4 µM of hemin and further red shift to 420 nm was observed upon further addition of hemin. (E) Fluorescence quenching studies of hemin titrated to FlrB-Fl^7-340^ shows its binding with saturation. K_d_ value 5.6 ± 0.02 μM obtained after fitting ΔF/ΔF_max_ versus hemin concentration. (F) AMP.PNP binding to FlrB-Fl^7-340^ is estimated through fluorescence quenching. Since saturation is not obtained the apparent K_d_ calculated was 172.03±0.025 μM. (G) Fluorescence quenching of FlrB-Fl^7-340^, incubated with hemin (4:1 molar ratio) upon addition of AMP.PNP showed better saturation with K_d_ of 105±0.024 μM. 3 mM hemin and 50 mM AMP.PNP stock was used for all fluorescence quenching experiment. (H) AMP.PNP binding to heme incubated FlrB-Fl^7-340^ was further checked through Isothermal Titration Calorimetry. The thermogram shows heat change along with lower panel of integrated isotherms. K_d_ value obtained was 158±10.5 μM after plotting non-linear regression to one side binding. All experiments performed in triplicate.

Fluorescence quenching experiments also indicated significant binding of hemin to FlrB-Fl^7-340^ with a K_d_ value of 5.6 ± 0.02 μM (Fig. 6E). Fluorescence quenching experiments conducted with AMP.PNP (λ_exc_=280 nm, λ_em_=295−400 nm) indicated binding, although saturation was not achieved. However, ΔF/ΔF_max_ vs AMP.PNP concentration plot yielded an apparent K_d_ of 172.03 ± 0.025 (μM) (Fig. 6F). This demonstrates approximately 30-fold higher efficacy of FlrB-Fl^7-340^ for heme, compared to AMP.PNP. Interestingly, similar quenching experiments between AMP.PNP, and hemin incubated FlrB-Fl^7-340^ (with hemin:protein molar ratio of 4:1), showed improved saturation with K_d_ of 105 ± 0.024 μM (Fig 6G), qualitatively demonstrating approximately two fold higher affinity for ATP upon heme incubation (Fig. 6F-G). Importantly, in this experiment, partial saturation of FlrB-Fl^7-340^ with hemin resulted significant reduction in initial fluorescence of FlrB-Fl^7-340^ (Fig. S8F, G). Complete saturation with hemin was drastically reducing overall fluorescence of FlrB-Fl^7-340^ whereTrp63 belonging to the globular region of PAS domain is the primary contributor to the overall fluorescence of FlrB-Fl^7-340^. Therefore, further quenching of fluorescence upon AMP.PNP binding to the CA domain, indicated that the conformational changes in CA domain upon AMP.PNP binding influences PAS domain conformation, plausibly because of their proximity in the 3D structure. We have validated the influence of hemin on ANP.PNP binding by ITC. While titration of AMP.PNP with free FlrB-Fl^7-340^ did not show any significant binding (Fig. S9), titration of hemin incubated FlrB-Fl^7-340^ with AMP.PNP, produced K_d_ of 158 ± 10.5 µM (Fig. 6H). Results of fluorescence quenching and ITC thus specified improved binding of AMP.PNP to FlrB-Fl^7-340^ upon hemin incubation.

### *V. cholerae* showed enhanced swimming motility with the increase in heme uptake

Heme is utilized as vital source of iron by most of the Gram-negative bacteria for their growth inside the host system. It acquires iron from host and utilizes it for its own cellular processes (Richard et al., 2019). As we mentioned before, enhanced expressions of the flagellar genes increase motility of *V. cholerae*. Phosphorylation of the bEBP FlrC by FlrB is a vital step in flagellar synthesis and our current study depicted that heme acts as a stimulus by binding to the PAS domain of FlrB. We have therefore investigated the effect of hemin on swimming motility of *V. cholerae* by a soft agar motility assay using an already established protocols (Luo et al., 2016; Martinez et al., 2009). *V. cholerae* was freshly grown from overnight culture till it reached O.D_600_ of ∼0.6. 5 µl culture was poured at the middle of each plate containing increasing concentration of hemin up to 200 µM. Plates were kept at 310 K for 12h, 24h and 48h. Continuous increase in zone of motility was observed with increase in hemin concentration (Fig. 7A-D). As hemin was dissolved in DMSO, motility assays were performed with plates containing equivalent amount of DMSO as control. The plate containing highest concentration of DMSO has no effect on motility since the zone obtained here is equivalent to the control plate with no heme (Fig 7A, E). Diameters of motility zones, calculated for each of the hemin concentration were plotted against respective incubation times using Origin8.5 (Fig. 7F). Maximum and consistent increase in motility was noticed with 200 µM of hemin.

**Figure 7:**
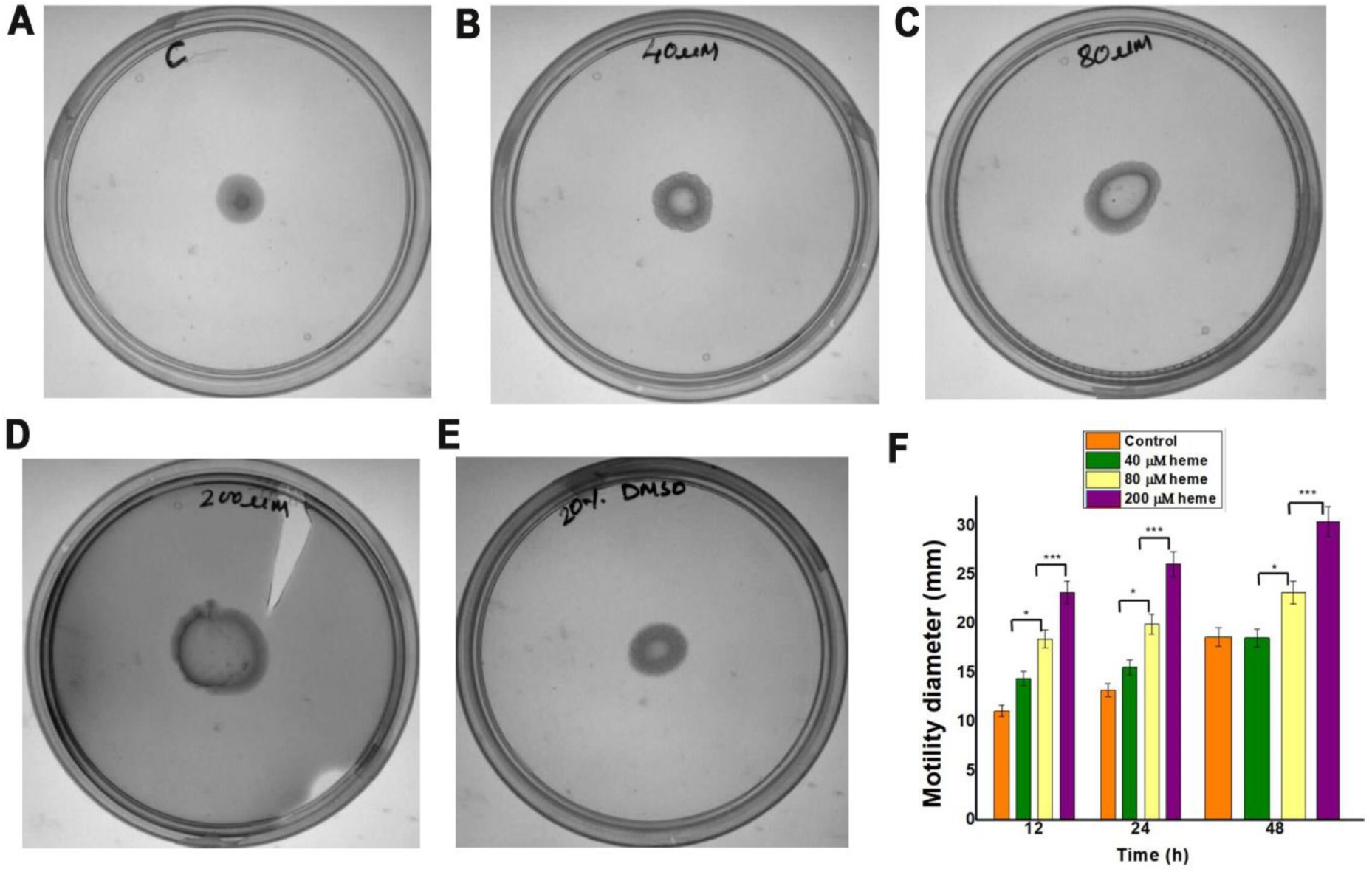
Motility of *V. cholerae* increases in the presence of hemin. Data collected after 12h, 24h and 48 h. Here we have presented the motilities after 24h incubation. (A) Control plate with no heme added, shows motility but with small zone. (B) Zone of motility gradually increased with (B) 40 µM (C) 80 µM (D) 200 µM of hemin. (E) Zone of motility of highest concentration of DMSO (diameter of motility zone is similar to that of control plate). Experiments were performed in biological triplicates. (F) Diameters of motility zones, calculated for each of the heme concentration along with respective incubation times are plotted here. Keeping time constant, significant increase in motility has been observed with increase in hemin concentration. Experiments were performed at least in triplicate mean ± SD. Two-way ANOVA was performed along with Tukey correction to compare each condition. * P < 0.05, *** P < 0.001.

### Sequence analysis indicated conservation of the heme binding residues

Apart from *V. cholerae*, we have identified 23 other monotrichous *Vibrio* species which possess FlrB and/or similar HKs. Although PAS domains are well known for sequence diversities, multiple sequence alignment depicted significant extent of identities among the PAS, DHp and CA domains of these *Vibrio* species (Fig. 8A). Such remarkably high sequence identities of the PAS domains of FlrBs highlight their architectural and functional similarities. Interestingly, we identified similar PAS domain containing histidine kinases in motile bacterial species like *Psychrobacter sp, Shewanella algae* etc. through careful BLAST searches with different search criteria. Multiple sequence alignment with those sequences revealed that the hydrophobic residues that constitute the ligand binding cleft of FlrB PAS (Fig. 1H) are mostly conserved in all those bacterial species (Fig. 8B). But, Tyr64, the potential iron coordinating residue located at the ligand binding groove, is not conserved. Rather, this has been replaced by different hydrophobic or basic residues (Fig. 8B). This further points towards heme binding in PAS through hydrophobic packing, rather than iron coordination.

**Figure 8:**
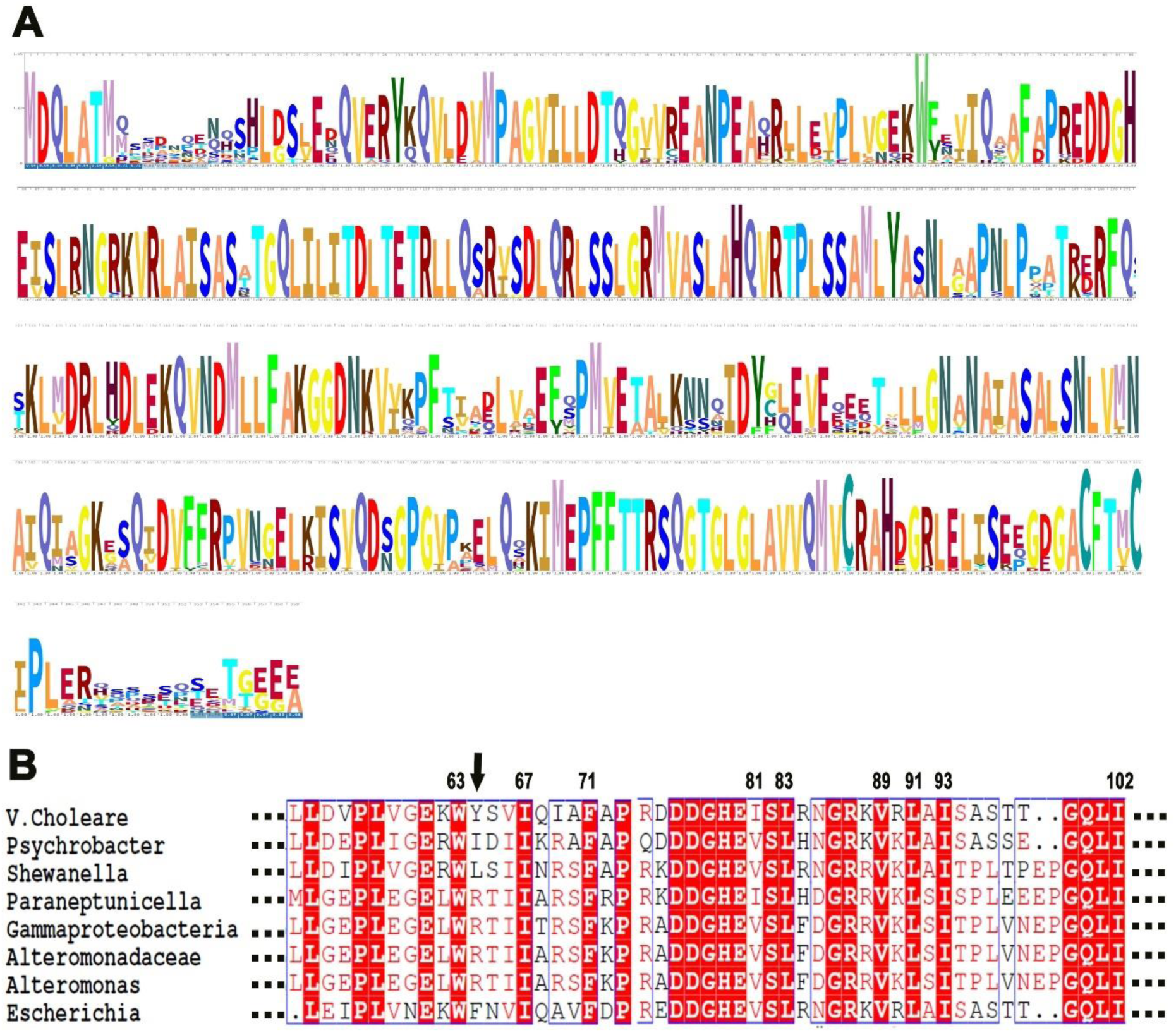
Hydrophobic residue involved in heme binding are conserved. (A) *Vibrio* species were searched for FlrB and/or similar HKs. Sequence alignment of FlrB of 24 *Vibrio* species including *V. cholerae such as V. tubiashii, V. galatheae, V. nereis, V. scophthalmi, V. ponticus, V. xiamenensis, V. mimicus, V. furnissii, V. fluvialis, V. gazogenes, V. aerogenes, V. diazotrophicus, V. anguillarum, V. qinghaiensis, V. campbellii, V. harveyi, V. alginolyticus, V. azureus, V. rotiferianus, V. tasmaniensis, V. mediterranei, V. nigripulchritudo and V. tapetis* which possess two component system FlrBC for flagellar synthesis depicted high extent of sequence identity. (B) Hydrophobic residues involved in the formation of heme binding pocket in FlrB-PAS are found to be conserved in similar kinases of motile bacteria while Tyr64 (shown by arrow) was not conserved.

## Discussion

Modular architecture of the histidine kinases (HKs) along with cellular localization of signalling domains are the pivotal factors to control their functions. In case of HKs, diversities in the signal recognition and transduction modules such as PAS or HAMP are more common, compared to DHp and CA domains. PAS domains execute a plethora of functions within sensory proteins by promoting protein-protein interaction (Lee et al., 2008; Ma et al., 2008; Neiditch et al., 2006), signal transfer (Oka et al., 2008), or by directly sensing perceived stimuli (Jonathan T. Henry1 and Sean Crosson1, 2012). In addition, plasticity of the dimeric interface of PAS often leads to diverse form of PAS dimers (Martinez et al., 2009). Although, FlrB/FlrC has been identified as unique TCS in controlling flagellar synthesis of *V. cholerae,* sensory signal and molecular basis of function of the cytosolic HK FlrB were elusive yet. The monomer of FlrB-PAS^7-123^ exhibits novel architecture and forms a stable parallel dimer involving Aα’, Hα and N-terminal loop (Fig. 1, 2, 4). Although the dimerization pattern of FlrB-PAS^7-123^ is largely similar to NifL of *Azotobacter vinelandii*, Aer2 of *P. aeruginosa*, or DosH of *E. coli* (Airola et al., 2013; Key et al., 2007; Park et al., 2004) distinct inter-domain interactions between extended N-terminal loop of one subunit with the globular region of the other contribute significantly to the robustness of FlrB-PAS dimers (Fig. 1C). So much so that FlrB-PAS^7-108^, that lacks Hα helices, also exhibits dimeric structure (Fig.4B, D).

Since FlrB is a cytosolic kinase, it is logical to assume that the molecule that enter cytosol would act as potential sensory ligand. Heme is an essential source to uptake iron by pathogenic bacteria for their growth and survival. Import of heme across the outer membrane of bacteria is accomplished by a TonB-dependent system with the help of proton motive force. In *V. cholerae*, active transport of heme across the plasma membrane is achieved by specific ATP binding cassette (ABC) transporters such as HutCD-B. The evolutionary pressure of producing numerous biochemical functions, together with the versatile chemical nature of heme, has given rise to different structural associations between heme and heme-binding proteins (Schneider et al., 2007). The question of how heme is attached to heme-binding proteins, depends on the shape and environment of the heme binding pocket. Existing results suggested that PAS domains of the sensory HKs usually bind heme tightly inside the cleft made of β-sheet and the helix Fα (Fig. 2D). However, a transient interaction demands a rapid association and dissociation of heme in order to allow for a situation-dependent response (Karnaukhova et al., 2014; Schneider et al., 2007) To the best of our knowledge, our current study provides the first report of heme binding to a sensory HK involved in flagellar synthesis. Since regulation of flagellar synthesis requires fine balance between motility and colonization, tight binding of heme might not be logical here. Hydrophobic interactions and π-π stacking of porphyrin ring, as well as electrostatic interactions and hydrogen bonds via the propionate sidechains were observed before in several heme binding proteins (Schneider et al., 2007). The globular region of FlrB-PAS is of unique architecture with a shallow cleft, and not a deep groove. However, its well-defined hydrophobic region, made of apolar residues arranged in layers (Fig. 1H), is suitable to pack the porphyrin ring. Furthermore, no clear reduction in hemin binding in the mutants Y64G, H79N or Y64A-H79A suggests the absence of the coordination of Tyr64 or His79 with heme iron. Rather, prominent binding of heme to FlrB-PAS and aforesaid mutants points towards heme binding through hydrophobic interactions. Ligand docking by ProBis further emphasized hydrophobic packing of heme with FlrB without metal coordination (Fig. S3). Notably, heme binding through hydrophobic interactions without iron coordination were recently witnessed in the other heme binding proteins as well (Gao et al., 2018; Tan et al., 2013). As for example, haemophore-like protein, HusA of *Porphyromonas gingivalis* was proven to bind heme exclusively through hydrophobic packing since no reduction in heme binding was observed upon mutation of the probable heme iron coordinating Tyr residue (Gao et al., 2018).

Trp63, located in the globular region of FlrB-PAS, is the major contributor to the intrinsic fluorescence of FlrB (Fig. 5K). Fluorescence quenching of FlrB-Fl^7-340^ upon AMP.PNP binding in the CA domain, therefore, indicates proximity of the CA and PAS domains in 3D structure of FlrB. Spectral changes during heme binding to Cytochrome P450 with peak around 425 nm and trough at 390–405 nm was considered as heme binding in low spin state (Horie & Watanabe, 2002; RenéFeyereisen, n.d.). In this regard, red shift of Soret band to λ_max_ of 420 nm with a trough at 360-390 nm, upon hemin titration with AMP.PNP treated FlrB-Fl^7-340^ (Fig. 6D) might be considered as synergistic mode of heme and ATP binding to FlrB. The binding affinity of heme towards FlrB-Fl^7-340^ is ∼30 times higher compared to that of ATP (Fig. 6E, F). Therefore, in a pull of heme and ATP in cytosol, heme is expected to get the priority to bind FlrB. We hypothesise that hydrophobic packing of heme to the PAS domain acts as a conformational switch to trigger efficient binding of ATP to the CA domain of FlrB. Notably, a heme binding domain dependent regulation of an ATP-dependent potassium channel was observed in a eukaryotic system as well (Burton et al., 2016).

In *V. cholerae*, phosphorylation of the RR domain of FlrC by FlrB is pivotal for flagellar synthesis and motility. Motility assay of *V. cholerae*, performed by us, demonstrated consistent increase in its swimming motility with increasing heme uptake (Fig. 7). This prompted us to conclude that heme binding to the PAS domain of FlrB initiates a conformational signalling that ultimately facilitates phosphorylation of FlrC. Eventually that triggers transcription of class-III flagellar genes leading to enhanced swimming motility of *V. cholerae.* Such regulation might be operational for FlrBC systems of similar *Vibrio* species, and acts as a prototype for flagellar synthesis in similar pathogens.

## Experimental Procedures

### Cloning, site-directed mutagenesis, overexpression and purification

FlrB-Fl^7-340^ (aa 7-340; Accession no. A0A0H3AMQ9) was cloned in pET-21b(+) vector using genomic DNA of *Vibrio cholerae* O395 as template. FlrB-PAS^7-108^ and FlrB-PAS^7-123^ were subsequently cloned in pET-28a(+) between *Nde1* and *BamH1* restriction digestion sites. The clones were selected appropriately using *Escherichia coli* XL1-Blue cells with ampicillin resistance for FlrB-Fl^7-340^, and kanamycin resistance for FlrB-PAS^7-108^ and FlrB-PAS^7-123^. Selected clones were confirmed by restriction digestion and commercial DNA sequencing. FlrB-Fl^7-340^ was expressed with C-terminus 6×His tag. FlrB-PAS^7-108^ and FlrB-PAS^7-123^ were expressed with N-terminal 6×His tag, which was subsequently removed by biotinylated thrombin followed by the incubation with streptavidin agarose for biochemical studies. Mutant clones of Y64G, H79N and W63A were prepared by two-step PCR amplification method using FlrB-PAS^7-123^ plasmid as template. Amplicons were cloned and selected using the same protocol used for wild type FlrB-PAS^7-123^. Clones were confirmed by restriction digestion check and commercial sequencing. Double mutant Y64A-H79A was purchased from Biotech Desk Pvt. Ltd. This construct was cloned in pET-28a(+) vector between *Nde1* and *BamH1* restriction endonuclease sites. Received plasmid was further overexpressed into BL21 (DE3) cells similar to other mutants mentioned above.

In order to purify the protein, single colony of the construct was inoculated into 5 ml of LB broth and kept in shaking condition overnight at 310 K. 500 ml LB was inoculated with the overnight culture and grown at 310 K until the O.D_600_ was of 0.6. At this stage, cells were induced with 1 mM Isopropyl ß-D-1-thiogalactopyranoside (IPTG) and kept under shaking condition for 3 hrs. Induced culture was then harvested at 4500g for 20 mins and resuspended in 5 ml of ice-cold lysis buffer containing 50 mM Tris-HCl (pH 8), 300 mM NaCl, 10 % Glycerol (v/v). 1mM PMSF (Phenylmethylsulphonyl fluoride) and 1 mg ml^-1^ lysozyme was added to the resuspended solution and lysed by sonication on ice. The cell lysate was then centrifuged at 12000 RPM for 50 mins at 277 K. Supernatant was collected immediately and poured onto Ni^2+^–NTA (Qiagen) resin (equilibrated with lysis buffer). 6×His-tag containing protein gets immobilized onto the pre-equilibrated Ni^2+^–NTA resins following the principal of immobilized metal affinity chromatography. Impurities were removed washing the resin with imidazole concentration 10 mM to 30 mM and the protein of interest was eluted with imidazole gradient 70 mM to 150 mM. Imidazole was removed from collected fractions through buffer exchange (50 mM Tris-HCl pH 8, 300 mM NaCl, 10% glycerol) using Amicon Ultra centrifugation unit (of molecular-weight cut off 3 kDa and 10 kDa) and further concentrated. Protein homogeneity was checked using 10% and 12% SDS PAGE. FlrB-PAS^7-123^ and FlrB-Fl^7-340^ were also purified in the presence of hemin. For the experiment, 300 uM of hemin was added to the lysate and and the proteins were purified as per protocol used for all other constructs and mutants. Ni-NTA purified fractions were checked through UV-vis spectroscopic scan. Purity of construct was checked using 10% and 12% SDS PAGE for FlrB-Fl^7-340^ and FlrB-PAS^7-123^ respectively.

### Crystallization

Crystallization trials were conducted for all three constructs. Crystals of FlrB-PAS^7-108^ were obtained by hanging drop vapour diffusion method at 293 K using 24 well crystallization tray with Ammonium Sulfate Grid Screen from Hampton Research with protein of concentration 10 mg ml^-1^. Mountable crystals were obtained when 2 µl of protein was incubated with 1.5 µl of precipitant containing 0.8 M Ammonium sulfate, 0.1 M Citric Acid pH 5, 10% Glycerol against 500 µl of the screening solution containing 2.4 M Ammonium sulfate, incubated at 293 K for 48 hrs. Crystallization of FlrB-PAS^7-123^ was also carried out using hanging drop vapour diffusion method as stated before with protein of concentration 10 mg ml^-1^. For FlrB-PAS^7-123^, initial crystallization trials were made using Molecular Dimensions SG1 Screen. Tiny plate shaped crystals were obtained with the precipitant containing 0.2 M Sodium acetate trihydrate, 25% (w/v) PEG 3350, 0.1M Sodium HEPES pH 7.5 against reservoir containing 500 µl of screening solution. Condition was optimized further and mountable crystals were finally obtained when 3 µl of protein and 2 µl of precipitant containing 0.2 M Sodium acetate trihydrate, 25% (w/v) PEG 3350, 0.1M Sodium HEPES pH 7.5, 17% (v/v) ethylene glycol was incubated against the solution with same contents without ethylene glycol in the reservoir at 293 K, for 72 hours.

### X-ray diffraction data collection and processing

The Crystals were soaked in the cryoprotectant having 30% (v/v) ethylene glycol in mother liquor with 0.5 mm litho loops of Molecular Dimensions, and flash frozen in liquid nitrogen. Diffraction data sets were collected at ID23-1 beamline of ESRF, Grenoble France at 100 K. Best data statistics were obtained by EDNA (Gordon & Bowler, 2013) processing software at ESRF. The data were then truncated using *AIMLESS* CCP4 (Winn et al., 2011).

### Phasing and model refinement

While trying molecular replacements with different structures of PAS domain no acceptable solution was found. In some cases, only four molecules were obtained with unacceptable packing and low LLG (e.g., with 4GCZ Top LLG = 75.5 and Top TFZ=6.9). Preparation of Se-Met derivative was not wise because the construct contains only one Met. Therefore, along with the preparation of the heavy atom derivatives of the crystals, we prepared a theoretical model of PAS construct (aa 7-123) using AlphaFold. Molecular replacement calculations by Phaser-MR of PHENIX (Adams et al., 2010) with monomeric AlphaFold2 generated model as search template using data between 48.2 and 2.4 Å resolution identified 12 monomers in the asymmetric unit with Top LLG = 1041.17 and Top TFZ=13.9 in space group C2. Symmetry consideration using graphical display indicated existence of six dimers of FlrB-PAS^7-123^ in the asymmetric unit. Initial rigid body, positional and B-factor refinement were carried out using PHENIX (Adams et al., 2010), which resulted a model with R_work_ of 37.4% and R_free_ of 41.7%. 2F_o_-F_c_ map calculated with this refined model was of good quality and overlaid with the model. Data collection parameters and refinement statistics are given in Table 1.

### Ligand Docking

For Docking, AutoDock Vina (OLEG TROTT, 2009) was installed along with MGLTools. Protein PDB and ligand PDB were first converted to pdbqt files. Single docking command was given through CMD to get 9 best binding modes. Output file was analysed using PyMOL as well as Discover studio client app.

### Spectroscopic titration assays

To observe possible Soret band shift of hemin upon binding with different FlrB constructs and FlrB-PAS mutants, 10 µM of purified protein was subjected to hemin (3 mM) titration through UV-vis spectroscopy. Hemin was added to the protein up to 46 µM successively as per requirement. After addition of each aliquot, sample was incubated for 10 mins and scanned between 250 to 700 nm. Final spectral shift was obtained after subtraction of free hemin spectra from hemin-protein spectra.

### Native PAGE

Purified 6 × His-tagged FlrB-PAS^7-123^ (200 µM) in buffer containing 50mM Tris-HCl (pH 8), 300 mM NaCl, 10% Glycerol was incubated with escalating concentration of hemin, from 400 µM to 1.2 mM for 45 mins at 293 K and NATIVE PAGE (12%) was conducted at 160 V as per protocol described in (Agarwal et al., 2017). Hemin stock of 5 mM was freshly prepared in 0.1 N NaOH. For control, protein was kept at 293 K for 45 mins without hemin. Experiment was performed in triplicate. Mobility shift of PAS domain was observed from 600 µM of hemin.

### Fluorescence quenching

Fluorescence measurement was carried out using a spectrofluorometer, Hitachi F-7000 protocol described by (Chakraborty et al., 2020). Changes in fluorescence of tryptophan and tyrosine were measured at an excitation wavelength of 280 nm and emission spectra were recorded between 295 nm and 400 nm with slit width of 5 nm for both excitation and emission wavelength. All reactions were carried out at 298 K. For all experiments, we used buffer containing 50 mM Tris-HCl (pH 8), 300 mM NaCl, 10% Glycerol. We used two ligands hemin (Sigma) and AMP.PNP. Hemin was at first dissolved in 100% DMSO to prepare 3 mM stock. This hemin stock was used for quenching study of FlrB-Fl^7-340^, FlrB-PAS^7-108^, FlrB-PAS^7-123^ and its mutants W63A, Y64G, H79N, Y64A-H79A. 50 mM AMP.PNP stock was used for quenching study of FlrB-Fl^7-340^ as well as for hemin incubated FlrB-Fl^7-340^. 5 µM of purified protein was subjected to fluorescence quenching with increasing ligand concentration starting from zero.

The dissociation constant, Kd was determined using nonlinear curve fitting analysis as per Equations 1 and 2. All experimental points for the binding isotherms were fitted by the least squares methods.

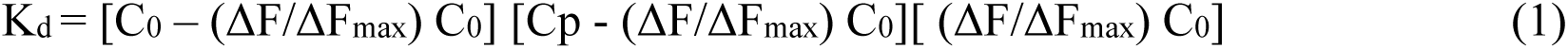

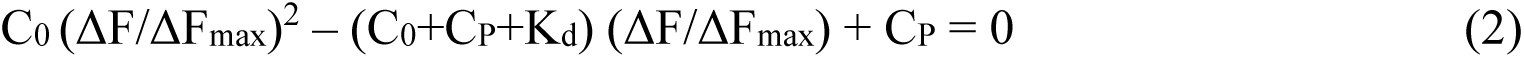

C_0_ denotes input concentration of ligands whereas C_p_ denotes the same for all protein construct used during experiment. ΔF is the change in fluorescence intensity at 323 nm (λ_ex_=280nm) for each point of titration curve and ΔF_max_ is the same parameter when ligand is totally bound to the protein. A double-reciprocal plot of 1/ΔF against 1/(Cp−C_0_), as shown in equation [3] was used to determine the ΔF_max_.

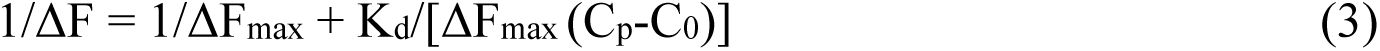

ΔF_max_ was calculated from the slope of the best-fit line corresponding to the above plot. All experimental data points of the binding isotherm were fitted by linear fit analysis using Origin 8.5 software.

### Isothermal titration calorimetry

For ITC experiment, Calorimetric measurement was performed using Nano ITC Isothermal Titration Calorimeter (TA Instruments, USA). Cell volume used is 350 µl of hemin sample and syringe was filled with 100 µl of equilibrated protein. Fresh PAS^7-123^, FlrB-Fl^7-340^ protein was prepared and dialysed with buffer containing 50 mM Tris-HCl (pH 8), 150 mM NaCl, 0.4% DMSO. 5 mM hemin stock was prepared in 100% DMSO and then diluted with buffer containing 50 mM Tris-HCl (pH 8), 150 mM NaCl to prepare working stock of 20 µM. 20 µM heme was titrated with various concentration of PAS^7-123^. In order to nullify the heat of dilution caused by DMSO, each heme Protein titration graph was prepared after subtracting titration of protein with buffer (50 mM Tris-HCl, pH 8.0, 150 mM NaCl, 0.4% DMSO).

Since the ligand solution contained DMSO, reverse titrations were carried out keeping ligand concentration (20 µM) in the cell and the protein in the syringe (400 µM). Both protein and heme solutions were equilibrated at 25°C and extensively degassed before loading to the cell or syringe prior to the titrations. Binding was carried out at 25°C with 2.5 µL of injection volume for each titration at stirring speed of 75 rpm. 20 injections each of 2.5 µl was applied at intervals of 180 s to 300 s. Peak integration and calculation of binding parameters were performed with Origin8.5 software by fitting to a one site binding model.

### Size exclusion chromatography

Dimerization of FlrB-PAS^7-123^ and FlrB-PAS^7-108^ was investigated through size exclusion chromatography. Gel filtration experiment were carried out at room temperature with Superdex 200 Increase 10/300 GL column from AKTA purifer (Cytiva). The column was first calibrated with standard molecular weight calibration kit (GE Healthcare) made of blue dextran containing Ferritin (440 kDa), Aldolase (160 kDa), Ovalbumin (45 kDa) and Lysozyme (14 kDa). Column was equilibrated with buffer containing 50 mM Tris-HCl (pH 8), 150 mM NaCl. FlrB-PAS^7-108^ and FlrB-PAS^7-123^ of concentration 100 µM, 250 µM and 500 µM were injected and eluted with the same buffer used for equilibration. This experiment was conducted separately for each of the concentration for both the constructs. Eluted protein was analysed through SDS PAGE. The standard graph was prepared against relative elution volume (V_e_/V_o_) in X-axis [where V_e_ is the elution volume and V_o_ is the void volume] and the log molecular weight in Y-axis.

### Crosslinking experiments

For this experiment, FlrB-PAS^7-108^ and FlrB-PAS^7-123^ were purified with buffer containing 50 mM Phosphate (pH 8.0), 150 mM NaCl and 10% glycerol. In each case, 2µl of 2mg/ml protein was incubated with increasing concentration of EGS [ethylene glycol bis (succinimidyl succinate), Thermo Scientific] and incubated for 30 mins in room temperature under dark condition. Protein samples were incubated with 100 µM, 250 µM, 500 µM, 700 µM, 1mM, 3mM of EGS and quenched with 1M Tris-HCl (pH 7.5). SDS PAGE analysis displayed presence of both monomer (14.8 kDa) as well as dimer band (around 30 kDa). In these experiments, control lane (Lane C) shows protein without EGS. As crosslinking experiment showed stable dimerization of PAS domains of FlrB, we further investigated if there is any role of heme upon its dimerization. Hence, we incubated FlrB-PAS^7-123^ with hemin concentrations of 10 µM, 20 µM, 50 µM, 100 µM, 150 µM and 200 µM for 10 mins on ice. Then 500 µM EGS was added to each of the reactions and incubated for 15 mins at room temperature under dark condition. In this case the control lane (Lane C) contains protein without hemin and EGS.

### Soft agar plate motility assay

For swimming motility test, soft agar plates were prepared using LB + 0.3% agar. Petri dishes of 90 mm were used for the assay. Hemin stock was prepared in DMSO and filter sterilized before use. Plates were prepared with hemin concentration of 40 µM, 80 µM and 200 µM. Same percentage of DMSO containing plates were also prepared for control experiments. *V. cholerae* culture was freshly grown from the overnight culture at 310 K up to the O.D_600_ of ∼ 0.6. 5 μl of this culture was poured at the middle of each of the plates and kept at 310 K incubators. Readings were taken at 12h, 24h and 48 h. All experiments were performed at least in triplicate. Diameter of motility zone was measured and calculated after each given time interval using ImageJ software (https://imagej.nih.gov/ij/) and data analysis was done using Origin8.5 software

### Data availability

Atomic coordinates and structure factors of FlrB-PAS^7-123^ have been deposited in the PDB ID 7YRT.

## Acknowledgments

We are extremely grateful to Dr. David Flot, the beamline scientist of ID23-1, ESRF, Grenoble, France for his tireless support during data collection. We are very thankful to Prof. Deepak T. Nair and Dr. Ramesh C. of RCB, India for their cordial support during sending crystals to ESRF, France. We are thankful to Prof. Udayaditya Sen of SINP, Kolkata, India for his constant support and valuable suggestions. We also thank Dr. K. Pal and Dr. T. Chakrabortty of Prof. U Sen’s lab at SINP, Kolkata for their help at different stages of this study. PM is thankful to her seniors Dr. Maitree Biswas, Dr. Sanjay Dey and the lab mates Shrestha Chakraborty, Indrila Saha, Ruchira Das and Arnab Pal for their constant help and support. We acknowledge Dr. Biplab Ghosh of RRCAT, Indore for his various suggestions. PM and JD are thankful to Dr. Sayak Ganguli of St. Xavier’s College (Autonomous), Kolkata for his valuable inputs in statistical analysis. We are grateful to Rev. Dr. Dominic Savio, SJ, Principal, St. Xavier’s College (Autonomous), Kolkata, for his constant support and encouragement. This work is supported by DBT, West Bengal grant No. 636(sanc)/BT(Estt)/RD-54/2015 and DST/INSPIRE Fellowship/2017/IF170190. We also acknowledge DST grant no. SR/FST/COLLEGE-014/2010(C), WBDBT BOOST grant no. 335/WBBDC/1P-2/2013 and DBT BUILDER grant no. BT/INF/22/SP41296/2020 for infrastructural support.

## Author contributions

JD conceptualized the study; PM, SA and SBM contributed to methodology; JD and PM contributed to writing—original draft preparation; JD, PM and SA contributed to writing, review and editing; JD supervised the study and acquired funding. All authors have read and agreed to the published version of the manuscript.

## Conflict of interest

We declare no competing financial interests.

## Notes

### Competing Interest Statement

The authors have declared no competing interest.

