## Supplementary Figures and Tables for "PAS domain of flagellar histidine kinase FlrB exhibits novel architecture, and binds Heme as sensory signal in unconventional fashion"

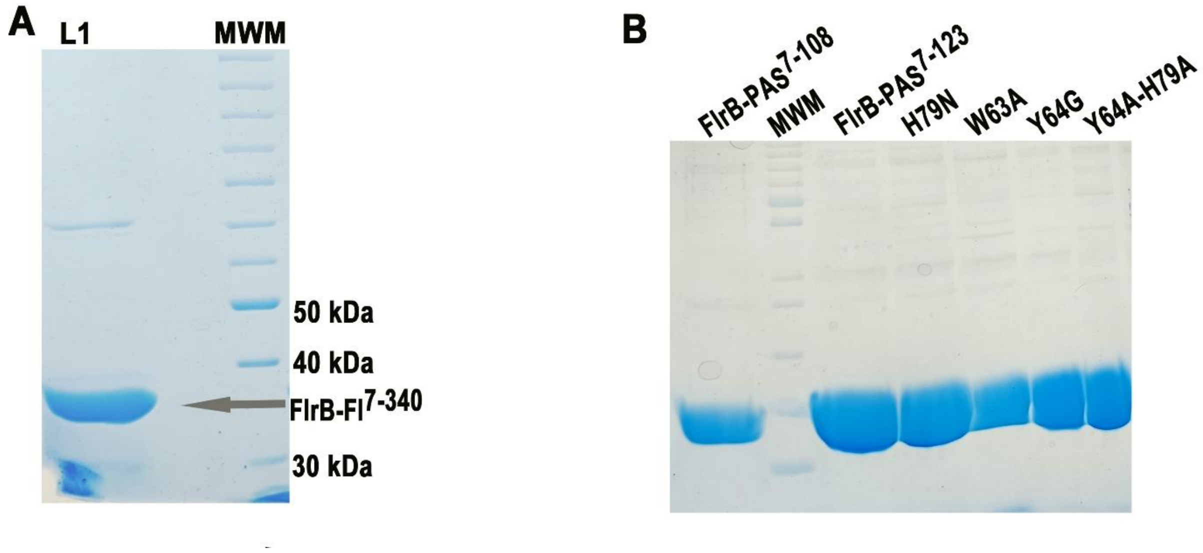


**Figure S1:** (A) 10% SDS-PAGE showing purification of FlrB-Fl$^{7\text{-}340}$. L1: 4µl of purified and concentrated FlrB-Fl$^{7\text{-}340}$ and next lane is the MWM. (B) 15% SDS-PAGE Purity of FlrB-PAS$^{7\text{-}108}$, FlrB-PAS$^{7\text{-}123}$ and the mutants of FlrB-PAS$^{7\text{-}123}$.

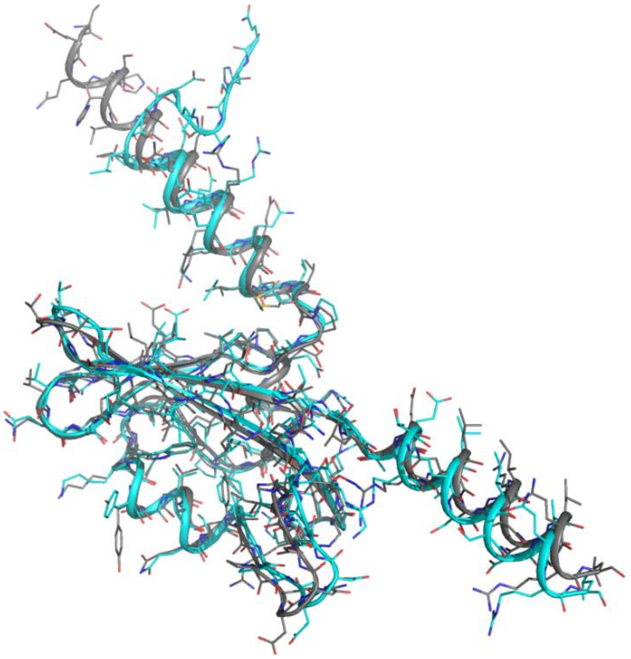

**Figure S2:** Superposition of the monomer of the crystal structure of FlrB-PAS$^{7\text{-}123}$ (cyan) on that of corresponding AlphaFold2 model (grey).

**Table S1: Protein structure comparison search in DALI server**

| No. | Chain | Z Score | RMSD | Lali (the number of structurally equivalent residues) | Nres (total number of amino acids in the hit protein) | %id (Percentage of identical amino acids over structurally equivalent residues.) |
|---|---|---|---|---|---|---|
| 1 | 5luv-B | 10.5 | 5.2 | 109 | 147 | 15 |
| 2 | 4gcz-A | 10.5 | 2.3 | 103 | 366 | 20 |
| 3 | 5fq1-A | 9.7 | 1.7 | 82 | 110 | 17 |
| 4 | 6gg9-D | 9.7 | 3.0 | 101 | 135 | 12 |
| 5 | 5j4e-C | 9.7 | 2.6 | 98 | 134 | 11 |
| 6 | 5j3w-C | 9.2 | 2.7 | 99 | 134 | 12 |
| 7 | 5fq1-B | 9.6 | 2.0 | 83 | 110 | 17 |
| 8 | 3sw1-A | 9.5 | 3.5 | 101 | 134 | 14 |
| 9 | 5xgb-A | 9.3 | 5.2 | 100 | 551 | 23 |
| 10 | 7a6p-A | 9.3 | 2.5 | 102 | 149 | 13 |
| 11 | 3lyx-B | 9.2 | 2.3 | 87 | 120 | 17 |
| 12 | 5xgd-A | 9.2 | 5.4 | 101 | 548 | 23 |
| 13 | 3b33-A | 9.1 | 2.4 | 82 | 109 | 20 |
| 14 | 3olo-A | 9.0 | 2.1 | 83 | 111 | 17 |
| 15 | 2r78-C | 9.0 | 2.4 | 87 | 116 | 16 |

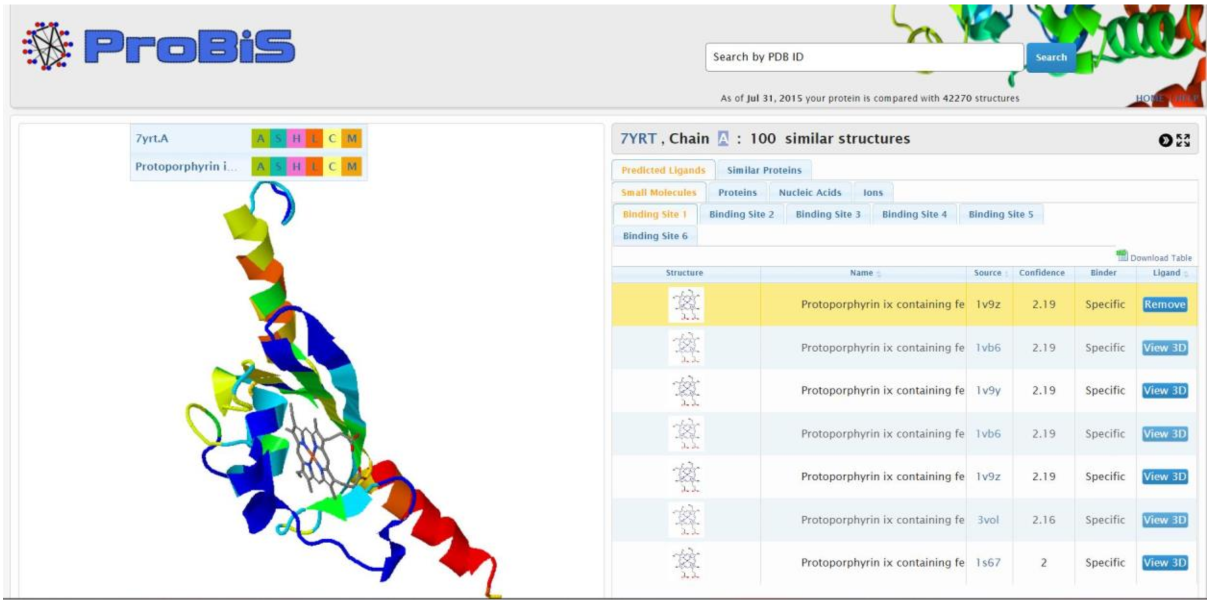


**Figure S3:** ProBis ligand prediction server identified Heme as ligand of FlrB-PAS$^{7-123}$

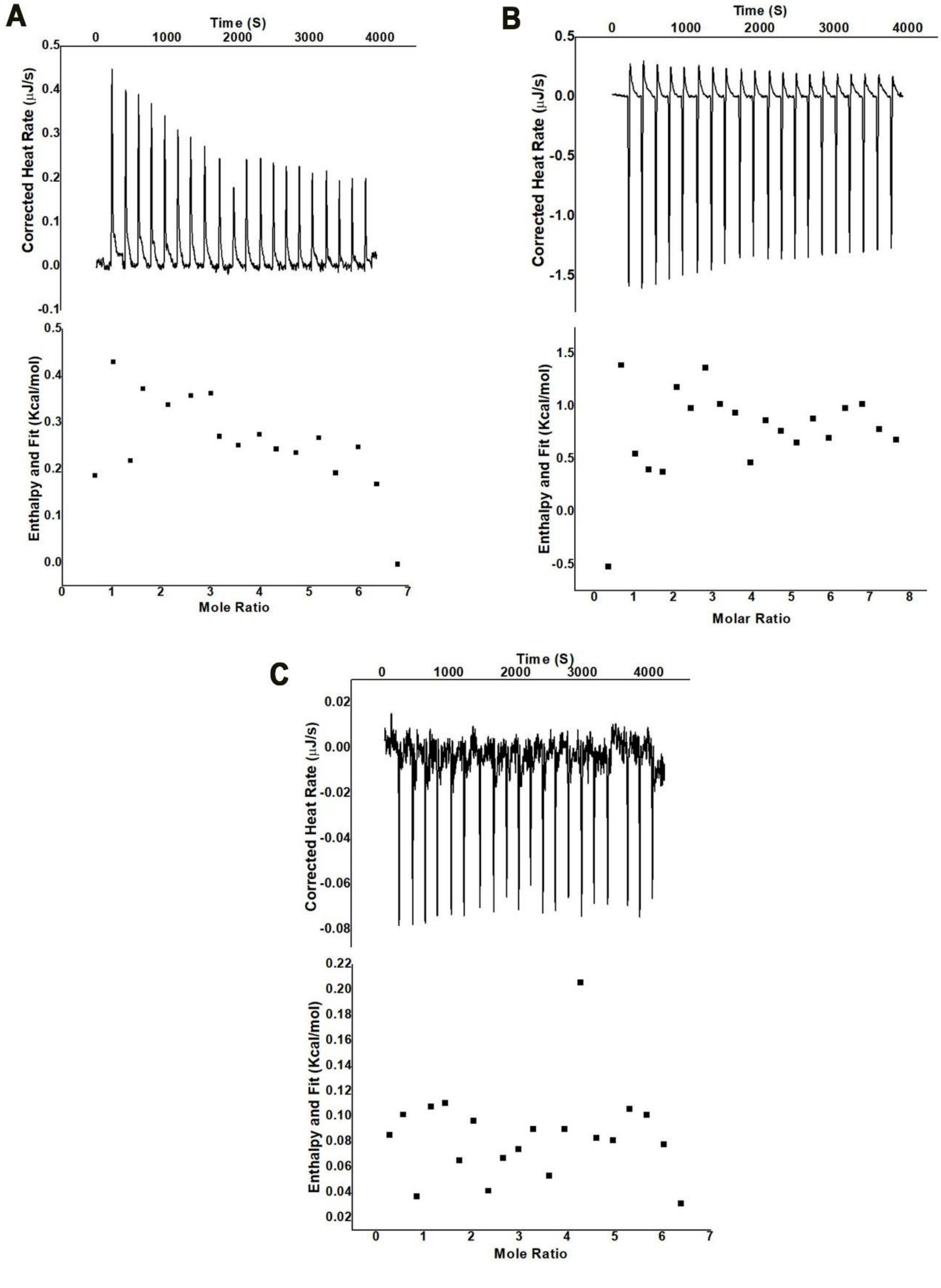


**Figure S4:** Upon buffer subtraction, isothermal Titration Calorimetry showed no significant binding of (A) FAD, (B) p-coumaric acid (C) DPO with FlrB-PAS$^{7\text{-}123}$.

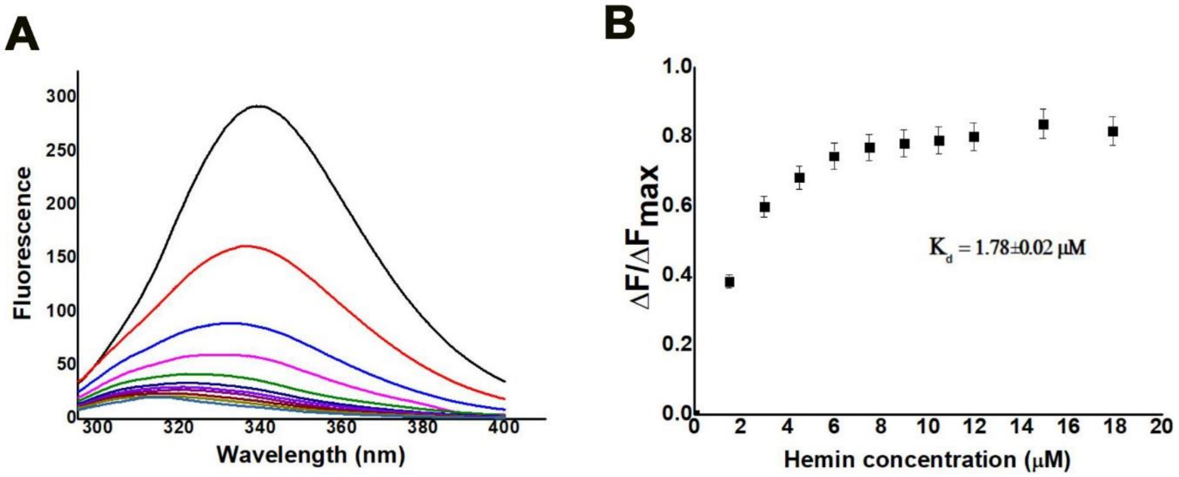


**Figure S5:** BSA is used as positive control to check the binding with hemin. (A) Binding of hemin with BSA has been tested through fluorescence quenching. (B) Plot of $\Delta F/\Delta F_{max}$ vs. Hemin concentration (μM) produced $K_d$ of 1.78 ± 0.02μM.

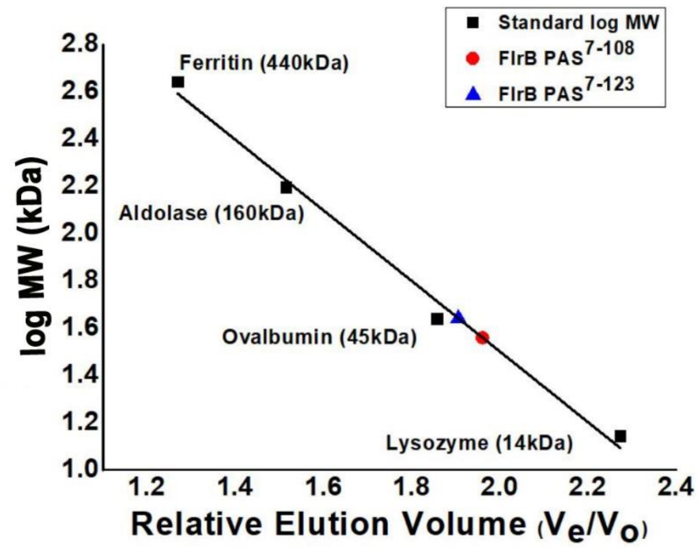


**Figure S6:** The calibration curve prepared using molecular weight standards by size exclusion chromatography in Superdex 200 Increase 10/300 GL column at AKTA purifier.

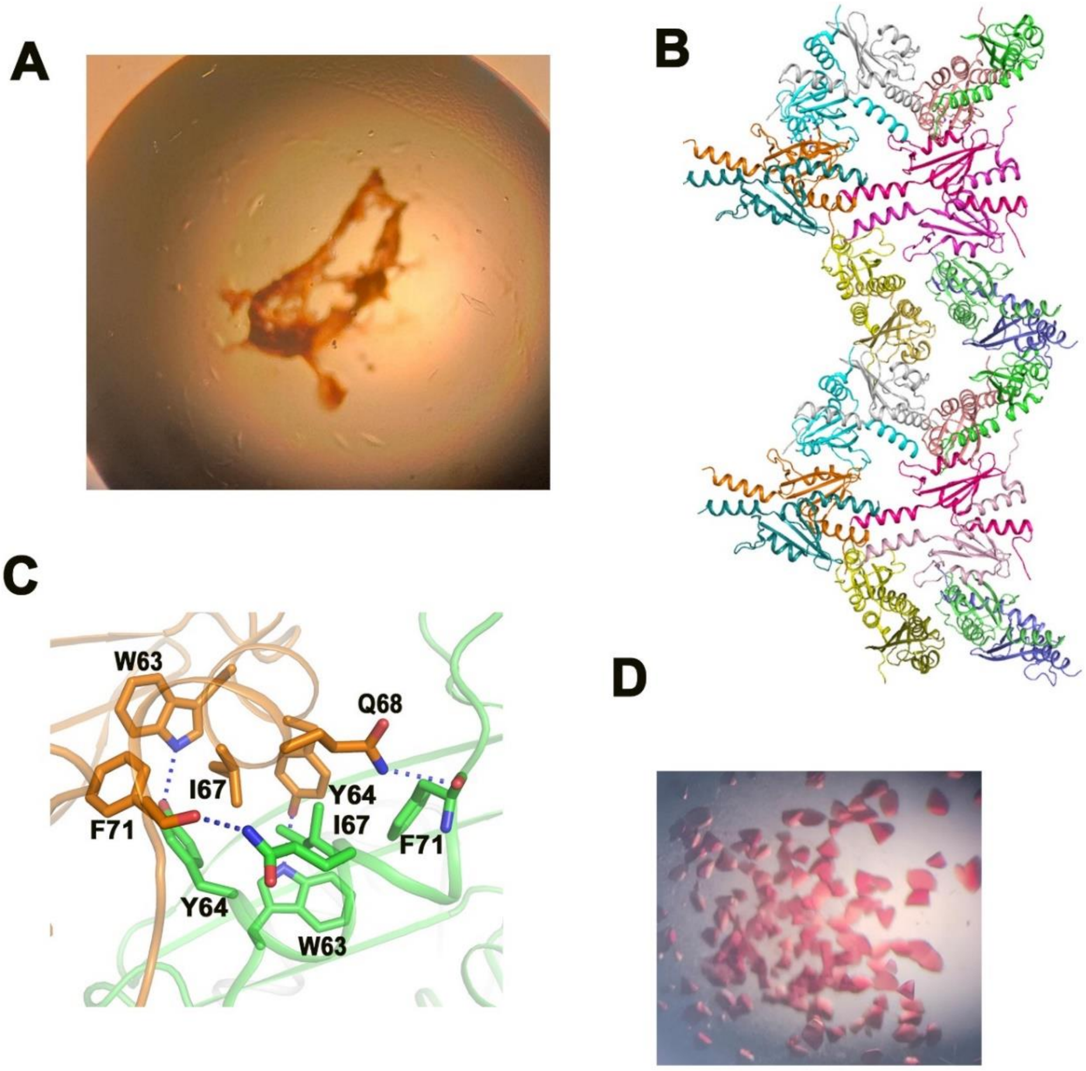


**Figure S7:** (A) Soaking of FlrB-PAS$^{7\text{-}123}$ crystals with hemin did not produce hemin bound crystal. (B) Packing of FlrB-PAS$^{7\text{-}123}$ dimers inside crystal. (C) Packing of the dimers inside crystal involves ligand binding cleft. Chain shown in brown belongs to one dimer and green is from another symmetry related dimer. (D) Hemin bound crystals of FlrB-PAS$^{7\text{-}108}$ diffracted very poorly.

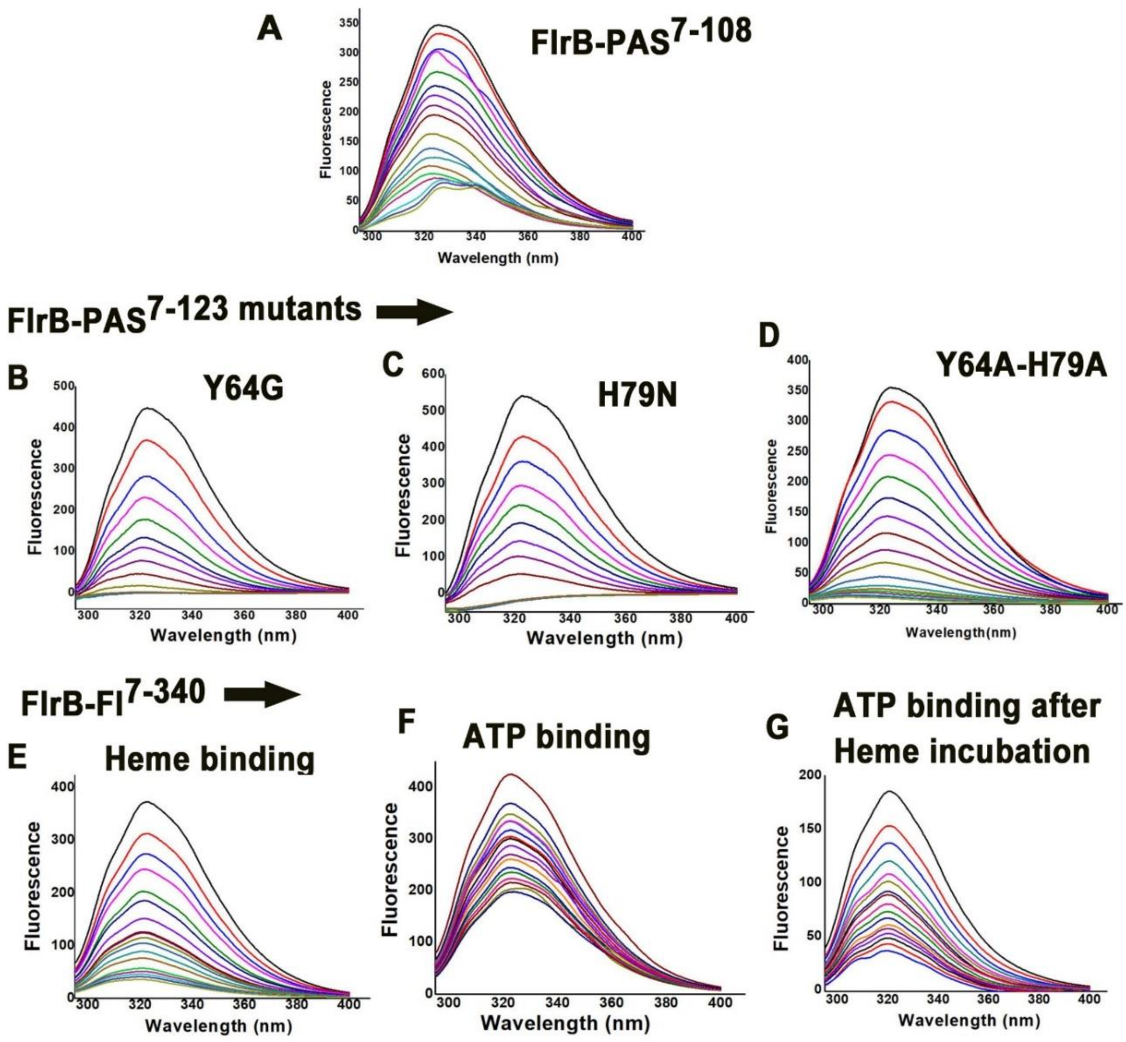


**Figure S8:** Fluorescence quenching graphs of (A) FlrB-PAS$^{7\text{-}108}$, (B-D) the mutants of FlrB-PAS$^{7\text{-}123}$ and of (E-G) FlrB-Fl$^{7\text{-}340}$ at different conditions.

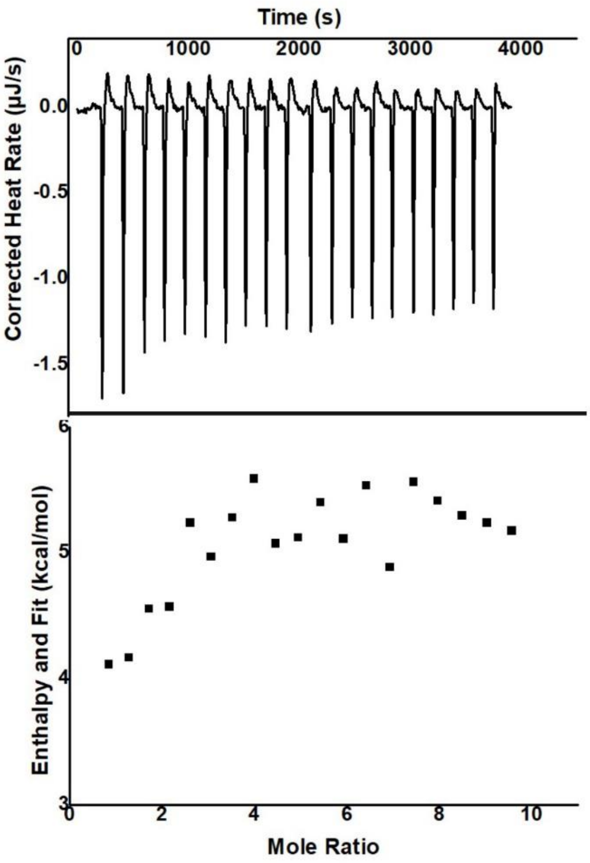


**Figure S9**: Isothermal Titration Calorimetry experiment to monitor AMP.PNP binding to FlrB-Fl$^{7\text{-}340}$. No measurable heat change was observed after buffer subtraction.
